# Deciphering Salt Stress Responses in *Solanum pimpinellifolium* through High-Throughput Phenotyping

**DOI:** 10.1101/2023.08.15.553433

**Authors:** Mitchell Morton, Gabriele Fiene, Hanin Ibrahim Ahmed, Elodie Rey, Michael Abrouk, Yoseline Angel, Kasper Johansen, Noha O. Saber, Yoann Malbeteau, Samir Al-Mashharawi, Matteo G. Ziliani, Bruno Aragon, Helena Oakey, Bettina Berger, Chris Brien, Simon G. Krattinger, Magdi A.A. Mousa, Matthew F. McCabe, Sónia Negrão, Mark Tester, Magdalena M. Julkowska

## Abstract

Soil salinity is a major environmental stressor affecting agricultural productivity worldwide. Understanding plant responses to salt stress is crucial for developing resilient crop varieties. Wild relatives of cultivated crops, such as wild tomato, *Solanum pimpinellifolium*, can serve as a useful resource to further expand the resilience potential of the cultivated germplasm, *S. lycopersicum*. In this study, we employed high-throughput phenotyping in the greenhouse and field conditions to explore salt stress responses of a *S. pimpinellifolium* diversity panel. Our study revealed extensive phenotypic variations in response to salt stress, with traits such as transpiration rate, shoot mass, and ion accumulation showing significant correlations with plant performance. We found that while transpiration was a key determinant of plant performance in the greenhouse, shoot mass strongly correlated with yield under field conditions. Conversely, ion accumulation was the least influential factor under greenhouse conditions. Through a Genome Wide Association Study, we identified candidate genes not previously associated with salt stress, highlighting the power of high-throughput phenotyping in uncovering novel aspects of plant stress responses. This study contributes to our understanding of salt stress tolerance in *S. pimpinellifolium* and lays the groundwork for further investigations into the genetic basis of these traits, ultimately informing breeding efforts for salinity tolerance in tomato and other crops.

## Introduction

Climate change and anthropogenic factors have exacerbated soil salinization, posing a significant challenge for agriculture in many regions (Qadir et al., 2014; Daliakopoulos et al., 2016; Tomaz et al., 2020; Eswar et al., 2021). Consequently, the development of salt-tolerant crop varieties is key for ensuring the stability of global food production. In the quest to future-proof agricultural systems, integrating salt tolerance mechanisms from non-domesticated species presents a viable approach compared to domesticated crops. Crop wild relatives evolved under diverse environmental conditions and thus exhibit a wide range of genetic variation for stress tolerance, including salt stress (Colmer et al., 2006; Sehrawat et al., 2013; Monteiro et al., 2018). The use of wild relatives has been implemented to improve abiotic and biotic resistance of cultivated high-value crops, such as tomato (*Solanum lycopersicum*) (Yassin, 1985; Ebert and Schafleitner, 2015; Torgeman and Zamir, 2023). *Solanum pimpinellifolium* is the closest relative to cultivated tomato (Pease et al., 2016), and inhabits diverse environments where species survival relies on substantial environmental resilience (Gibson and Moyle, 2020). Previously, *S. pimpinellifolium* was used as donor for introgression of multiple agronomic traits, ranging from pathogen resistance to fruit morphology (Yassin, 1985; Sarfatti et al., 1991; Paran and van der Knaap, 2007; Ashrafi et al., 2009; Celik et al., 2017; Zhang et al., 2018). The generation of high-quality reference genomes of *S. pimpinellifolium* (Razali et al., 2018; Wang et al., 2020a), enables the development of natural diversity panels and forward genetic studies.

Advances in imaging-based phenotyping allow for more precise dissection of plant responses to the environment. Phenotyping platforms that integrate various sensors facilitate simultaneous study of multiple traits throughout the stress exposure. The non-destructive nature of imaging-based phenotyping provides opportunities to study responses to stress at high spatial and temporal resolutions (Dhondt et al., 2013; Fiorani and Schurr, 2013; Tardieu et al., 2017). Additionally, automation and image processing allows evaluation of many genotypes. Indoor and outdoor phenotyping systems rely on different methods for image collection, and provide various spatial, spectral and temporal resolution (Fiorani and Schurr, 2013; Araus and Cairns, 2014; Tardieu et al., 2017). Measuring plant phenotypes at higher resolution and under controlled environmental conditions can provide a more accurate insight into trait variation and thus result in more precise heritability estimates (Houle et al., 2010; Tardieu et al., 2017; Ziliani et al., 2018; Roitsch et al., 2019). However, evaluating plant performance in field conditions, and validating the specific genotype performance in more diverse and agronomically realistic scenarios is also of high relevance for selecting specific germplasm for future breeding programs.

In this study, we developed a well-genotyped diversity panel of *S. pimpinellifolium,* which facilitates future studies on abiotic and biotic stress resilience present within this particular wild tomato species. We leveraged the power of high-throughput phenotyping to systematically assess the morphological and physiological responses to salt stress across *S. pimpinellifolium* accessions in greenhouse and field conditions. By revealing the intricate dynamics of salt stress response in this wild relative of the cultivated tomato, we aim to identify key genetic factors which could be harnessed to improve salt tolerance in agricultural tomato varieties and other crop species. This research represents a step forward in our understanding of plant salt stress responses, potentially contributing to the development of more sustainable and resilient agricultural systems in the face of growing environmental challenges.

## Materials & Methods

### Plant genetic material

In total, 274 *Solanum pimpinellifolium* accessions (**Table S1**) were used in this study. Individual genotypes were propagated for two generations in the KAUST greenhouses, with bees excluded from the greenhouse and plants kept physically separated to prevent cross-pollination. Additionally, the *S. lycopersicum* genotype (Heinz 1706) was included in the field experiment.

### DNA extraction

Nuclear DNA was isolated from desiccated leaf tissue using NucleoSpin Plant II kit from Macherey-Nagel, followed by the Ribonuclease A treatment included in the kit. The quality of the DNA was examined on 2% agarose gel and the quantity was estimated using nanoDrop. The DNA samples were shipped to NovoGene for whole genome sequencing, including quality control, preparation of 350 bp insert DNA library, and sequencing using Novaseq PE150 (**Table S1**).

### Read mapping and SNP calling

SNP variant calling was performed following the method described in (Abrouk et al., 2020) and available on GitHub (https://github.com/IBEXCluster/Wheat-SNPCaller) with a few modifications. Raw sequence reads were filtered with Trimmomatic-v0.38 (Bolger et al., 2014) using the following criteria: SLIDINGWINDOW:5:20; MINLEN:50. The filtered paired-end reads were then aligned for each sample individually against the LA2093_genome_v1.4 reference assembly (Wang et al., 2020a) using BWA-MEM (v-0.7.17) (Li and Durbin, 2010), only reads mapping with a quality Q>20 were retained, followed by sorting and indexing using samtools (v1.8). Duplicated reads were marked and read groups were assigned using the Picard tools (http://broadinstitute.github.io/picard/). Variants were identified with GATK (v4.1.8.0) (McKenna et al., 2010) using the “--emitRefConfidence’ function of the HaplotypeCaller algorithm and to call SNPs and InDels for each accession (Van der Auwera et al., 2013). Individual g.vcf files for each sample were then compressed and indexed with tabix (v-0.2.6) (Li, 2011) and combined into chromosome-chunks g.vcf using the GenomicsDBImport function of GATK. Joint genotyping was then performed for each chromosome-chunk using the function GenotypeGVCFs of GATK. To obtain high confidence variants, we excluded SNPs with the VariantFiltration function of GATK with the criteria: QD < 2.0; FS > 60.0; MQ < 40.0; MQRankSum < −12.5; ReadPosRankSum < - 8.0 and SOR > 3.0. Full chromosome VCF files and subsequently full genome VCF were obtained using the gatherVCF algorithm of GATK (Danecek et al., 2011). In total, 37,687,189 SNPs were called from 491 individuals (**Table S2**).

To obtain high-quality SNP data, we applied SNP clustering filter to allow no more than three SNPs in a 10-bp window using VariantFiltration from GATK v4.1.8.0 (argument: --cluster-size 3 -- cluster-window-size 10). Additional filters were applied with VCFtools (v 0.1.17) (Danecek et al., 2011) to obtain high-quality SNPs: low and high average SNP depth (6 ≤ DP ≥ 30), keep only bi-allelic sites, accepting missing data ≤10%, SNPs located in chromosome unanchored, and accessions having > 10% of missing data. Finally, 20,325,817 SNPs and 482 individuals were kept (**Table S2**). The SNP density was calculated in bin sizes of 100 kb using VCFtools (v0.1.17) and plotted using the ggplot2 R package (**Fig. S1**).

### Determining genetic structure within the population

We assessed the genetic relationships between individuals by principal component analysis (PCA) using all high-quality SNPs (20,325,817 SNPs) with PLINK software v1.90 (Purcell et al., 2007). We estimated individual ancestry coefficients using the sNMF function implemented in the R package LEA v2.0 (Frichot and François, 2015) with the entropy option and with 10 independent runs for each *K* (*K* is the number of putative ancestral populations) from *K=*1 to *K=*10. The analysis was run with randomly chosen samples of 120,000 SNPs. SNP density across the 12 chromosomes was calculated using VCFtools (v0.1.17).

### Characterization of salt stress-induced changes under controlled greenhouse conditions

The experiment was carried out at The Plant Accelerator, University of Adelaide, Australia, across two conveyor-based greenhouses - Northeast (NE) and Northwest (NW), each room fitted with 24 conveyor lanes and 26 carts per lane (see **Fig. S3**). The experiment used a split-plot design, with pairs of consecutive carts in lanes forming the main plots, both of which were planted with the same accession and randomly assigned either the control or salt treatment, with 288 main plots per greenhouse. The accessions were unequally replicated, with three replicates for 136 accessions and two replicates for 84 accessions. The allocation of accessions to main plots was determined using the DiGGer (Coombes) package in R v2.12.1 (Bunn and Korpela, 2014). One terminal cart per lane was left empty and another was filled with soil to measure soil evaporation, the evaporation pots being positioned systematically so that two occurred across the lanes in 13 of the 25 positions (see Figure S03).

The *S.pimpinellifolium* seeds were soaked in 50% household strength bleach for 25 min, rinsed twice in fresh water for 5 min before sowing at 1 cm depth, four seeds per pot. The seeds were sown in pots filled with a coco peat potting mix, containing ready-release and slow release fertilizer (MINI Osmocote 16-3-9+te). Each pot contained 1.43 kg potting mix and was maintained at a gravimetric water content of 42% (w/w). At 14 days after sowing, seedlings were thinned to one evenly sized seedling per pot. At 16 days after sowing pots were loaded onto the conveyor system and fitted with double-layered plastic mats to limit evaporation and distribute water more evenly from the automated watering system. Plastic trays were placed underneath each pot to retain water. From 17 days after sowing, the plants were imaged daily. And soil water content was maintained at 42% (w/w) by watering to target weight with rainwater. Salt stress was administered by adding 200 mL of a 0.6 M NaCl solution into the tray beneath each pot, to reach 200 mM NaCl in the soil solution once pots dried back down to the target weight. Control plants were treated with 200 mL of rainwater. Salt stress was applied at 21 and 24 days after sowing in NE and NW greenhouses respectively, corresponding to emergence of leaf 6, to account for slight developmental differences between the two greenhouses. During the salt stress application, the youngest emerging leaf was marked for identification at harvest for ion measurements.

The difference (*Δ*) in pot weight (*M*) between consecutive days (*t*) was used as an approximation of the volume of water transpired by the plant over that time (transpiration rate, TR), after correcting for evaporation from the soil surface by subtracting the weight difference observed in the nearest evaporation cart:

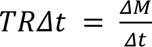

The LemnaTec RGB imaging station captured two side-view (SV1 and SV2) images at 90° rotation and one top-view (TV) image of each plant (Al-Tamimi et al., 2016). The sum of green pixels from these three views was used to calculate the projected shoot area (PSA):

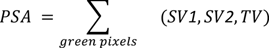

By measuring PSA over time, growth rates for individual plants can be calculated (Brien et al., 2020). Growth rates can be expressed in absolute terms as the increase of the PSA over time (Absolute Growth Rate, AGR), or in relative terms as the increase of the PSA over time as a proportion of the previous day’s PSA (Relative Growth Rate, RGR):

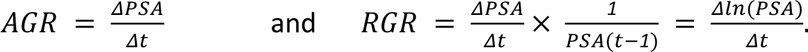

Additionally, by combining the daily transpiration rate with the increase in PSA, we can calculate the transpiration use efficiency (Al-Tamimi et al., 2016):

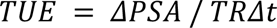

Plants were harvested 14 days after application of salt stress (35 and 38 days after sowing in NE and NW greenhouses, respectively). The shoot fresh mass was recorded for all harvested plants. Above ground biomass (except for leaves used for ion consent measurement) was passively dried at room temperature for one week, oven-dried at 80 °C for 24 h, and stored in plastic bags until measurement of shoot dry mass (SDM).

Measured shoot fresh mass was used to calculate a series of stress indices (**Fig. S6**): Salt tolerance index (STI1 = Salt / Control), Tolerance index (TOL = Control - Salt), Mean productivity index (MP = (Control + Salt) / 2), Geometric Mean Productivity (GMP = square root of (Control x Salt), Stress Susceptibility Index (SSI = ((Control - Salt) / Control)/ ((population mean Control - population mean Satl) / population mean Control), Stress Tolerance Index (STI2 = (Control x Salt) / population mean Control^2^), and Stress Weighted Performance Index (SWP = Salt / square root of Control).

### Ion content measurements of plants grown under controlled conditions

For quantification of leaf sodium (Na) and potassium (K) contents, the marked leaves that developed during the salt stress treatment were collected. We also collected a subset of control plants. The leaves were weighed for fresh weight, oven-dried for 24h at 80 °C, weighed for dry weight, and stored in a labeled 50 mL falcon tube. Subsequently, 20 mL of 1% nitric acid was added to each tube, incubated at room temperature for 24 h, followed by 24 h in the oven at 80 °C. The concentrations of Na^+^ and K^+^ ions in leachate dilutions were determined using a flame photometer (model 420; Sherwood Scientific Ltd., UK).

### Phenotypic data curation for plants grown under controlled conditions

The phenotypic data underwent manual data curation, to identify the plants that underwent considerable mechanical damage as they moved throughout the phenotyping facility due to their large size. The data was additionally inspected for abnormal changes in plant size over time, such as decrease in projected shoot area. The images of affected plants were inspected to see if abnormal data was caused by physical damage due to the size of the plants. In cases where plant damage was negligible, or if another explanation was found (e.g. the effects of salt stress, or a large, oddly oriented leaf that evaded capture by imaging), these minor anomalies were left unchanged.

Several errors in the automatic conveyor system led to changes in measuring schedules, affecting the PSA and transpiration measurements. Where changes to watering caused outlier values, transpiration data for such days were removed. On several occasions, some evaporation carts reported evaporation values vastly exceeding the water loss data of their assigned plant carts, leading to negative transpiration values. These erroneous measurements were replaced with the mean across all evaporation carts in the same greenhouse for that day.

### Processing of phenotypic data for plants grown under controlled greenhouse conditions

Cubic smoothing splines were used to characterize longitudinal traits over time (with 5 degrees of freedom), on account of their ability to describe plant growth dynamics without a priori assumptions on their shape (Brien et al., 2020). Based on the observations in general trends in plant growth dynamics, we have identified three intervals that allowed reduction of complexity of this dataset: interval 1 (day 0 to 5 after salt imposition), interval 2 (day 6 to 9 after salt imposition) and interval 3 (day 10 to 14 after salt imposition)Daily data were grouped by interval and summary statistics were calculated to provide an average measure for that time period. Spatial corrections were performed on data grouped by intervals in R using the ASReml-R package (Butler et al., 2009), which generated a Best Linear Unbiased Estimator-derived mean for each trait, by accession, interval and treatment. Data visualization was carried out in R, using the ggplot2 (Wickham, 2016), ggpubr (Kassambara), ggplotly (Sievert) packages. Trait-specific broad-sense heritability estimates were calculated using MVApp (Julkowska et al., 2019).

### Evaluation of *S. pimpinellifolium* lines under field conditions

199 *S. pimpinellifolium* accessions and one *S. lycopersicum* accession (Heinz 1706) were evaluated for their performance under field conditions (**Table S1**). The field site consisted of four 33 x 34.5 m fields (**Fig. S9**). The capacity of each field was 300 plants, arranged in 15 lanes of 20 plants. Plants were spaced 1.5 m along lanes and 2 m between lanes. A border surrounding each field was planted with Sudan grass and blue switch grass as a buffer for the edge effects. These specifications allowed for 600 experimental plants per treatment, i.e. 3 replicates for all 200 genotypes under each treatment (salt and control). An unbalanced replicated experimental design was calculated with DiGGer (Coombes) package in R v2.12.2 to randomize the positions of genotypes and treatment blocks across the field.

36-pot trays were filled with Metro-Mix 360 potting soil pre-mixed with fungicides at 2 mL/L and watered abundantly. We sowed 18 pots per genotype and 3 seeds per pot. Trays were watered, covered with a clear tray lid, and placed in the greenhouse at 28/20°C for day/night temperatures. After germination was observed (4-6^th^ October, 2017), the seedlings were grown at 23/20°C, and the lid was removed. Seedlings were thinned down to one per pot at the 2-leaf stage. Seedlings were watered, fertilized, treated with pesticides, and provided supplemental lighting as necessary. From October 26, day temperatures were increased by 1°C / day to reach 28°C on October 31, while night temperatures were maintained at 20°C. On October 29-30, 6 homogenous, representative seedlings were selected from each accession and arranged in trays reflecting the randomized design in which they would be planted in the field (Johansen et al., 2019b). All randomized trays were transported to the Agriculture Research Station of KAU at Hada Ash Sham (110 km north-east of Jeddah, altitude 226 m, 21°47ʹ48ʺN, 39°43ʹ35ʺE) on October 31 (Johansen et al., 2019b).

The soil type of the experimental sites was classified as sandy loam (sand 84%: silt 14%: clay 2%). The area’s climate is tropical arid with less than 100 mm annual precipitation. The experimental site was unused for 6 years, and prepared for the experiment by plowing the site in May 2017 to a depth of ∼ 50 cm using a tractor-drawn subsoiler, then plowed orthogonally using a disc plow, then leveled using a box blade. Cattle manure was applied at a rate of 10 tons/ha and monosuperphosphate (12.5% P2O5) at 150 kg/ha. The field was then left until September 2017. The experimental site was precisely leveled before installing the dripper lines on the soil surface along the pre-prepared rows. Dripper lines (Rain Bird model LD-06-12-1000, Saudi Arabia) were fitted with external drippers (4 L/h flow rate) every 1.5 m. Experimental plants were planted at each dripper such that irrigation water was delivered directly to their base. The downstream end of each dripper line was connected to a manifold leading to the water pumps. The inlet pressure on each dripper line was approximately 1.5 bars. The water source was from four holding tanks fed via the main irrigation network, which delivered low-salinity groundwater (∼30 mM NaCl) from a nearby borehole (see **Fig.S09B**).

Plants were transplanted into their assigned positions in the experimental field (November 1-2). Day and night temperatures ranged from 27 - 37°C and 12 - 24°C, respectively. The mean temperature throughout the growing season was 25.7°C. No rainfall was recorded over the course of the experiment. Several sand storms were observed on December 8 and 16, and January 4 and 8 to 10. Plants were rinsed following these events to remove dust from the plant canopies to avoid negative effects on plant physiology and image-based measurements. Both control plots were irrigated using the low salinity water groundwater source, as were the salt-treated plots prior to salt stress application. Based on preliminary field experiments, salt stress was applied incrementally over the course of the experiment to a final concentration of approximately 250 mM. Salt stress was first applied to salt-treated fields on November 14, by mixing a 3M stock solution of NaCl into the irrigation tanks of salt-treated fields to reach a final concentration of ∼200 mM. This method raised the salt concentration to ∼225 mM on December 10 and ∼250 mM on December 18. Due to additional stress caused by high winds, the salt concentration was reduced to ∼180 mM prior to harvesting on January 12. All fields were irrigated, with respective water sources, twice daily (morning and evening) for 10-minute periods from November 1 to 8, for 15 min periods from November 9 to December 16 and for 30 min periods after December 17.

### UAV phenotyping

UAV-based imagery was used to determine plant characteristics such as size, canopy temperature, and health status. Cameras mounted on UAVs were flown over the field site at a fixed speed and height following a pre-programmed flight path, capturing radiometric data at regular intervals.

A hyperspectral camera was used to assess plant area, height, volume, color, canopy cover, and condition (Johansen et al., 2019b; Johansen et al., 2020). TIR camera was used to capture canopy temperatures to inform the estimation of plant transpiration rates. RGB and TIR cameras were all mounted on a DJI Matrice 100 Quadcopter (Dà-Jiāng Innovations, China). The RGB camera used was a Zenmuse X3 camera (Dà-Jiāng Innovations, China), which captures radiometric data in the visible spectral range (400-700 nm), across 3 continuous bands (i.e., band center wavelengths R=605 nm, G=525 nm, B=460 nm). The TIR camera used was a ThermalCapture 2.0 640 thermal camera (<u>Malbéteau et al. 2021</u>), based upon a modified FLIR Tau 2 (TeAx, Germany). DJI Matrice 100 flights carrying these cameras were performed at approximately solar noon to reduce variation in light conditions and ambient temperatures and allow greater comparability of data between individual campaigns. Flights were carried out at a fixed height of 13 m and a constant speed of 2m/s. AgiSoft PhotoScan software (Agisoft LLC, Russia) was used to process RGB imagery producing georeferenced orthomosaics and a Digital Surface Model (DSM) for each campaign. A final resolution of 0.5 cm/pixel for RGB images was achieved. Radiometric calibration converted digital data from UAV-based imagery into at-surface reflectances. These image processing steps are described in detail in (Johansen et al., 2019a). The TIR camera had a 640 x 512 pixels resolution and a 13 mm focal length, collecting thermal infrared images across the 7.5-13.5 µm spectral range. The thermal data presented is limited to campaign dates that overlapped with the hyperspectral measurements, which determined the plant delineations required to extract data for each plant (Angel and McCabe, 2022). Due to a malfunction of the weather station used to calibrate the thermal imagery on November 30, thermal data is not available for this campaign.

A hyperspectral camera was used to generate narrowband and broadband vegetation indices (VIs) that inform on various plant characteristics and aspects of plant condition and explore methods to remotely determine pigment contents (see imagery processing and modeling details in (Angel and McCabe, 2022)). Hyperspectral imagery was collected using a DJI Matrice 600 (M600) hexacopter coupled with a Ronin-MX gimbal to reduce flight dynamic effects. The flight platform housed a Headwall (Headwall, USA) Nano-Hyperspec push-broom camera, with 12 mm lens and a horizontal field of view of 21.1°, which gathered radiometric data in the 400- 1000 nm spectral range across 272 continuous bands, with 6 nm full-width half maximum. An image co-registration approach was used to process hyperspectral imagery and georeferenced mosaics, achieving a spatial resolution of 0.7 cm/pixel for each campaign (see processing details in (Angel et al., 2019)). The empirical radiometric calibration method converted radiance data from UAV-based imagery into at-surface reflectances (see method details in (Barreto et al., 2019)). To estimate the Projected Shoot Area (PSA) based on the hyperspectral time series, a minimum noise fraction (MNF) transformation was applied to produce a new version of the images, whose bands are descending ordered by signal-to-noise ratio. Then, using the first three MNF-bands that keep the most contrast between plant canopy and soil background, an unsupervised classification approach was employed to delineate the pixels representing plant tissues. Reflectance data was extracted from all pixels assigned to each plant on each date – the mean reflectance values at different wavelengths across all pixels are used as the reflectance data for each plant. Two of the narrowband greenness VIs produced based on the hyperspectral dataset (**Table S7**) are considered in this study (see the full list of VIs in (Angel and McCabe, 2022)). The Modified Red Edge Normalized Difference Vegetation Index (MRENDVI) is a chlorophyll index that offers insight into plant health by assessing plant greenness using near-infrared and/or red-edge wavelengths highly reflected by green leaf tissues (Sims and Gamon, 2002) The Photochemical Reflectance Index (PRI) is also considered for its ability to assess photosynthetic light use efficiency, in particular by measuring reflectances in the yellow bands that inform on the xanthophyll class of carotenoid pigments (Sims and Gamon, 2002).

Extensive ground-truth validation was performed to enable the calibration and validation of UAV-derived data (**Tables S8 and S9**). Geometric correction of RGB, multispectral and hyperspectral imagery was achieved using five Ground Control Points (GCPs) deployed at each of the four corners and at the center of the field. Their GPS coordinates were measured at the time of transplanting (November 2nd) using a Leica GS10 base station with an AS10 antenna and a Leica GD15 smart antenna as a rover (Leica Geosystems, Switzerland) (see details in (Angel et al., 2019)). Radiometric calibration of RGB, multispectral and hyperspectral imagery was achieved using six radiometric calibration panels of painted plywood boards, whose reflectance values were measured at each campaign using a FieldSpec4 spectrometer (Malvern Panalytical, United Kingdom) (see details in (Angel and McCabe, 2022)). Direct measurement of several plant characteristics in the field was performed on a selected subset of plants in order to calibrate and validate UAV-derived data (**Table S9**). Six accessions were chosen on the basis of varying levels of salt tolerance in terms of biomass based on preliminary field trial results (two low, two medium, and two high tolerance). In addition to whole plant measurements, some measurements were carried out on selected leaves from reference plants. The three youngest fully expanded leaves on a single shoot were selected at the beginning of each campaign.

### Destructive phenotyping of field-grown plants at terminal harvest

Plants were harvested between January 18 and 26 to determine shoot fresh mass (SFM) and the number of mature and immature fruit (MFN: Mature Fruit Number; IFN: Immature Fruit Number) and their total mass (MFY: Mature Fruit Yield; IFY: Immature Fruit Yield). These data were used to determine the Total Yield Number (TYN, i.e., the total number of mature and immature fruit) and Total Yield Mass (TYM, i.e., the total mass of mature and immature fruit), the Mature Fruit Mass and Immature Fruit Mass (MFM and IFM, i.e. the average mass of mature and immature fruit, respectively) and the Harvest Index (HI, i.e. the harvestable fraction of above-ground biomass) (**Table S10**). Plants were cut at ground level and weighed to determine SFM. For plants with SFM < 1 kg, fruit measurements were taken from the whole shoot. For plants with SFM > 1 kg, fruit measurements were taken from a representative subset of the shoot (between ∼0.5 - 1 kg). All fruit larger than 3 mm was picked, and separated into immature and mature fruit. For plants for which fruit measurements were taken from a representative subset of the shoot, whole plant immature and mature fruit numbers and mass were extrapolated by multiplying the subset measurement by the ratio of the whole SFM by the mass of the representative subset.

Measured shoot fresh mass was used to calculate a series of stress indices (**Fig. S6**): Salt tolerance index (STI1 = Salt / Control), Tolerance index (TOL = Control - Salt), Mean productivity index (MP = (Control + Salt) / 2), Geometric Mean Productivity (GMP = square root of (Control x Salt), Stress Susceptibility Index (SSI = ((Control - Salt) / Control)/ ((population mean Control - population mean Satl) / population mean Control), Stress Tolerance Index (STI2 = (Control x Salt) / population mean Control^2^), and Stress Weighted Performance Index (SWP = Salt / square root of Control).

### Processing of phenotypic data for plants grown under field conditions

All data analysis shown here, including calculation of means and salt tolerance indices, correlation analyses, statistical tests and preparation of figures, was carried out in R (R Core Team, 2014). Broad-sense heritability estimates were calculated using MVApp (Julkowska et al., 2019).

### Further SNP filtering and GWAS analysis

We used the file ‘hard_filtered_snps_SPIMP.allchro.vcf’ which contains a total of 49,993,627 SNPs across 491 sequencing samples, as a starting point for the selection of SNPs for GWAS analysis. We removed all “SRX” samples, that indicated the accessions sequenced in other studies and thus not part of our collection. We used Beagle (version 18May20.d20 (Pook et al., 2020)) to impute the missing SNPs. Subsequently, we used bcftools (V.1.9) to remove all heterozygous SNPs. We removed LA1258, LA1572, LA1581, LA1280, LA1343, LA1576, LA1593, LA1633, LA1634, LA1685, LA2839, LA0859, LA3158, LA3161, and LA2659 from the file as two or more sequencing files represented them. SNPs with Minor Allele Count below three were removed using vcftools (Danecek et al., 2011). We used TASSEL-5 (Glaubitz et al.) to transform the vcf file format to a HapMap format. The HapMap file was used to run the Genome-Wide Association Studies using GAPIT (Wang and Zhang, 2021), as well as calculating the kinship matrix (**Fig. S19**). The SNP information was recalculated into a numerical matrix for the ASReml GWAS scripts (Korte et al., 2012). The kinship matrix, calculated using GAPIT, was included in the GWAS models.

### Data availability

The raw sequencing data of the 265 *Solanum pimpinellifolium* genotypes sequenced in this study are available on EBI-ENA under the study number PRJEB64210. The raw and filtered VCF files used for the population structure and diversity study are available on the DRYAD database https://datadryad.org/stash/share/96wpRaXlL2XNkbyMaJmVj241I2gulCRgRgqWh2Vizzg. The input data for the GWAS can be found at https://doi.org/10.5281/zenodo.7551780. The output data from the GWAS for the shoot data can be accessed at https://doi.org/10.5281/zenodo.5874309, whereas the R-notebook containing the analysis pipeline to identify the most interesting associations for the TPA data can be accessed at https://rpubs.com/mjulkowska/pimpinellifolium_GWAS_TPA.

## Results

### Genetic diversity within selected *S.pimpinellifolium* panel

To evaluate the genetic diversity that is specific to *Solanum pimpinellifolium*, we developed a panel of 274 *S. pimpinellifolium* accessions, with their place of origin spanning a wide range of habitats. Out of these, 265 accessions were sequenced, with genome coverage varying between 11 and 27 times the 0.9 GB estimated genome size (**Table S1**). The sequences were aligned to *S. pimpinellifolium* genome LA2093 (Wang et al., 2020a) together with 226 accessions of wild and cultivated tomatoes previously resequenced (Gao et al., 2019), and single nucleotide polymorphisms were called using GATK (McKenna et al., 2010). After filtering the SNPs, we identified 20,325,817 SNPs (**Table S2**), with the highest SNP number present in chromosome 1, and the lowest SNP number present in chromosome 6 (**Fig. S1**). The SNP density distribution across the chromosomes shows a lower SNP density towards the chromosome ends (**Fig. S1**).

To evaluate the variation within *S. pimpinellifolium* in comparison to cultivated tomatoes, we used publicly available data from NCBI of samples belonging to cultivated tomatoes. We performed the structure and diversity analyses on a total of 482 accessions. A diversity analysis using principal component analysis (PCA) demonstrated that *Solanum lycopersicum var. cerasiforme* exhibited the smallest cluster size and was grouped with *S. lycopersicum* (**Fig. 1 A**). *S. pimpinellifolium* and *S. lycopersicum* were found to be distributed along the PCA axis, forming distinct clusters (**Fig. 1 A**). We estimated the ancestry coefficient for each accession using all available samples **(Fig. S2)** and only the re-sequenced 265 *S. pimpinellifolium* samples (**Fig. 1 B-C**). In the *S. pimpinellifolium* population, the genetic clustering analysis revealed distinct groupings at different values of *K* (where *K* is the number of putative ancestral populations). At *K*=2, a few accessions were separated from the rest of the samples. At *K*=3, two small groups of accessions formed separate clusters. At *K*=4, the *S. pimpinellifolium* population was divided into four groups, with three clusters being distinct from the majority of samples and containing only a small number of samples. Additionally, we identified a small number of accessions (LA0859, LA3158, LA3160, LA3161, LA3159) that were likely misclassified as *S.pimpinellifolium*, as they are clustering with the *S. lycopersicum* (**Fig. S1**). These results suggest considerable genetic diversity within the *S. pimpinellifolium* population and the diversity gradient persistent over the North to South (i.e., latitude) axis.

**Figure 1.**
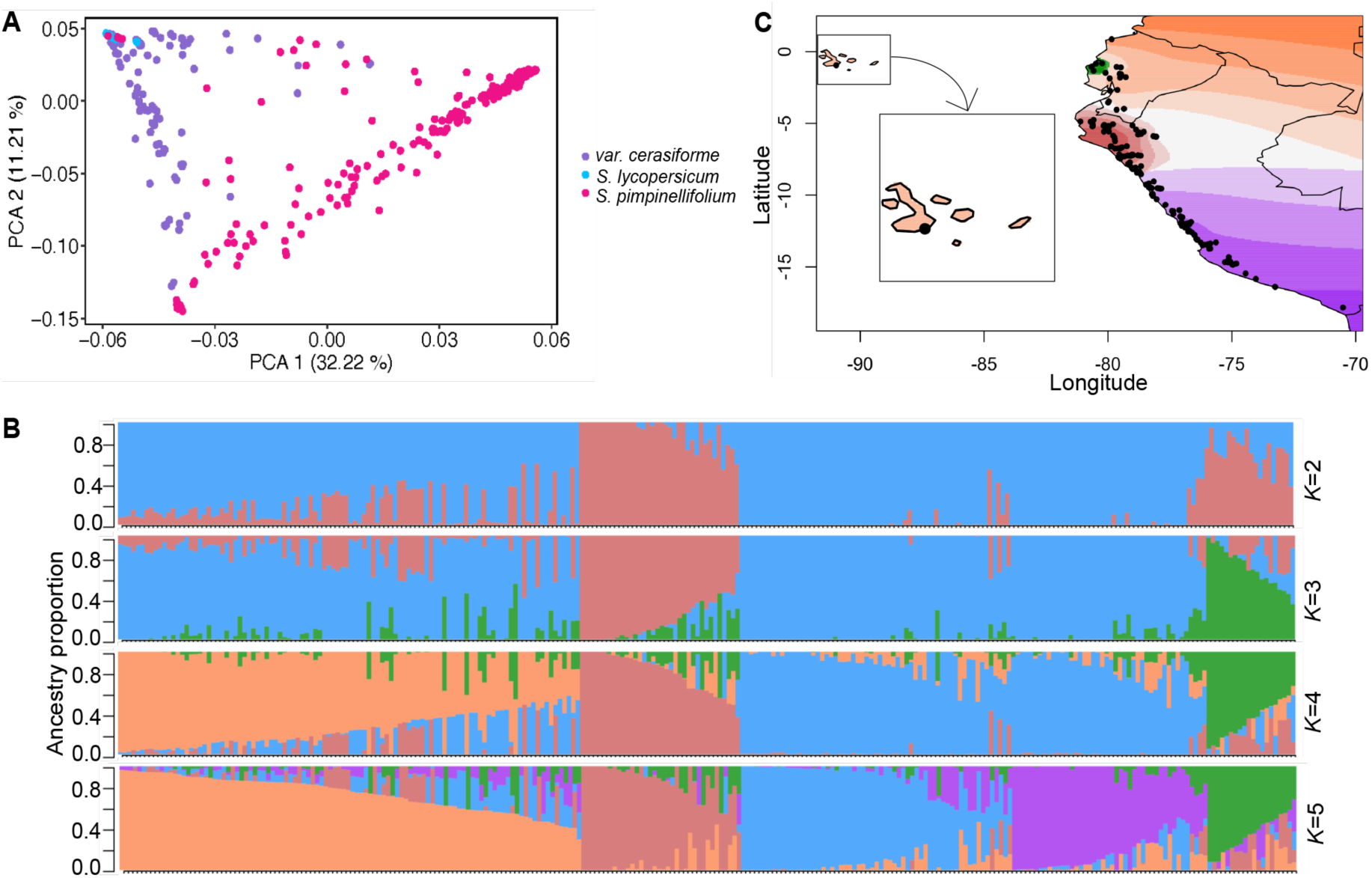
Genetic diversity and structure of *S. pimpinellifolium* and *S. lycopersicum*. **(A)** Principal component analysis (PCA) of 482 tomato samples using all SNPs (20,325,817 SNPs). In purple are 151 *S. lycopersicum var. cerasiforme* samples, in blue; 30 *S. lycopersicum* samples and 301 *S. pimpinellifolium* samples in pink. **(B)** Population structure (from *K* = 2 to *K* = 5) of 265 *S. pimpinellifolium* samples estimated with sNMF. Each bar represents a sample and the bars are filled by colors representing the likelihood of membership to each ancestry. **(C)** Geographic distribution of ancestry proportions of *S. pimpinellifolium* samples obtained from sNMF analysis at *K* = 5. The colors represent the maximal local contribution of an ancestry. Black dots represent the coordinates of each sample. Only samples with known GBS coordinates are shown (n=202).

### Salt tolerance of *S. pimpinellifolium* during early vegetative growth

To characterize the diversity in physiological responses to salt among the developed population, a total of 219 *S. pimpinellifolium* accessions were screened for salt-induced changes in shoot growth, transpiration rate, and ion accumulation under greenhouse conditions (**Fig. S3**). The seedlings were exposed to salt stress at six-leaf stage and phenotyped over two weeks, with daily RGB imaging, weighing, and watering. After 14 days of stress, the plants were harvested for destructive measurements, including shoot fresh mass and ion accumulation in the leaf that developed during the salt stress exposure. The strong correlation between the projected shoot area and shoot fresh mass (r^2^=0.931, **Fig. S4**) confirmed the use of projected shoot area as a reliable proxy for digital plant biomass.

The salt treatment induced observable differences in absolute and relative growth rates, as well as transpiration rate and transpiration use efficiency from the 1st day of treatment (**Fig. 2**). Throughout the experiment, absolute growth rate increased with a slowing trend towards the end of the imaging period. The absolute growth rate in salt-treated plants was significantly lower compared to control plants throughout the treatment period (**Fig. 2 A**). On the other hand, the relative growth rate gradually decreased over the duration of the experiment (**Fig. 2 B**), a pattern extensively described in annual crops (Hunt). Salt stress significantly reduced the relative growth rate at all observed time points (**Fig. 2 B**). While salt stress significantly decreased the transpiration rate throughout the experiment (**Fig. 2C**), it increased the transpiration use efficiency during the final phase of the experiment (**Fig. 2D**). To reduce the complexity of the dataset, we summarized growth rates, transpiration rate, and transpiration use efficiency over three intervals. The length of each interval was selected based on the patterns in the recorded data, with intervals 1, 2, and 3 taking place between days 0-5, 6-9, and 10-14 after salt stress treatment, respectively (**Fig. S5**). The patterns observed for each interval concisely reflected the continuous changes observed throughout the experiment.

**Figure 2.**
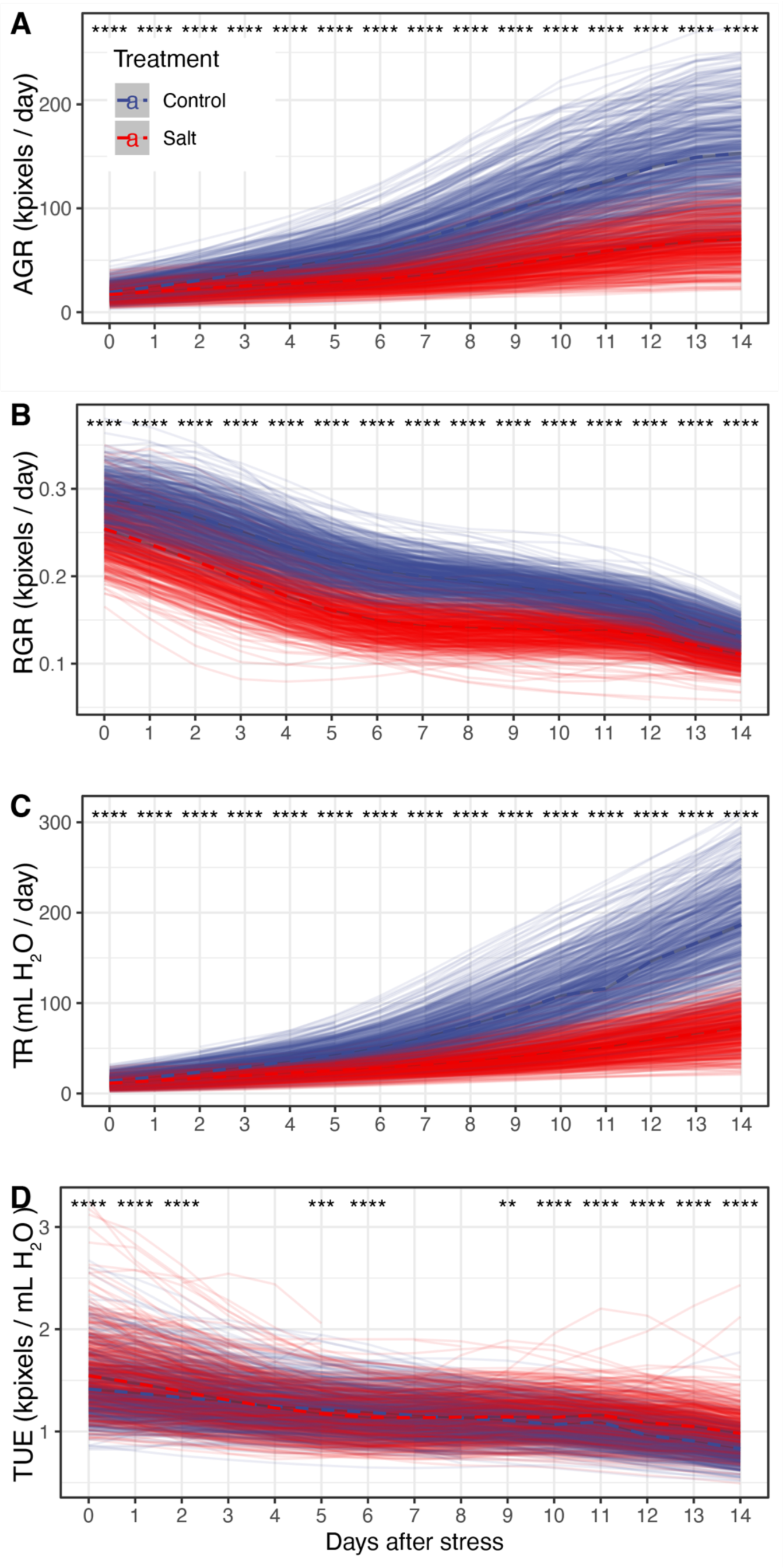
Salt stress reduced *S. pimpinellifolium* growth and transpiration. Natural diversity panel of *S. pimpinellifolium* was germinated in the greenhouse conditions and exposed to an effective concentration of 0 or 200 mM NaCl at 21-24 days after sowing, corresponding to Control and Salt stress treatments respectively. The plants were monitored for 2 weeks after treatment application for increase in the projected shoot area and daily transpiration rate. **A)** Absolute Growth Rate was calculated based on daily change in projected shoot area, while **B)** Relative Growth Rate was calculated using a proportion of increase in PSA to previou’s day PSA. **C)** Transpiration Rate was calculated by calculating the difference in pot mass between consecutive days after correcting for evaporation for soil surface as compared to the nearest empty pot. **D)** Transpiration Use Efficiency was calculated as a ratio of the projected shoot area and transpired water recorded for each day. The individual transparent lines describe the progression of individual experimental plants, while bold dashed lines describe population average. The differences between treatments were tested using one-way ANOVA, and *, **, *** and **** indicate p-values below 0.05, 0.01, 0.001 and 0.0001 respectively.

Terminally extracted traits revealed that salt stress treatment decreased fresh and dry shoot mass by 47% across the population (**Fig. 3 A-B**). While leaf Na^+^ content increased by 84% upon salt stress treatment (**Fig. 3C**), no significant difference was observed for the leaf K^+^ content (**Fig. 3C-D**). The broad sense heritability for all traits varied between 0.84 and 0.1 (**Table S4**). To examine the salt-specific effects, we calculated seven stress-tolerance indices using shoot fresh mass (**Fig. S6, Table S5**). Stress indices including Salt tolerance index (STI = Salt / Control), Tolerance index (TOL = Control - Salt), and Stress Susceptibility Index = ((Control - Salt) / Control)/ ((population mean Control - population mean Satl) / population mean Control)) highlight accessions that suffer the least under salt stress but do not necessarily perform well under non-stress conditions. On the other hand, Mean productivity index (MP = (Control + Salt) / 2), Geometric mean productivity (GMP = square root of (Control x Salt), Stress Tolerance Index (STI2 = (Control x Salt) / population mean Control^2^) and Stress Weighted Performance Index (SWP = Salt / square root of Control) indices identify accessions with high performance under both conditions, with little (STI2, GMP) or no (MP, GMP) consideration for the relative differences between them.

**Figure 3.**
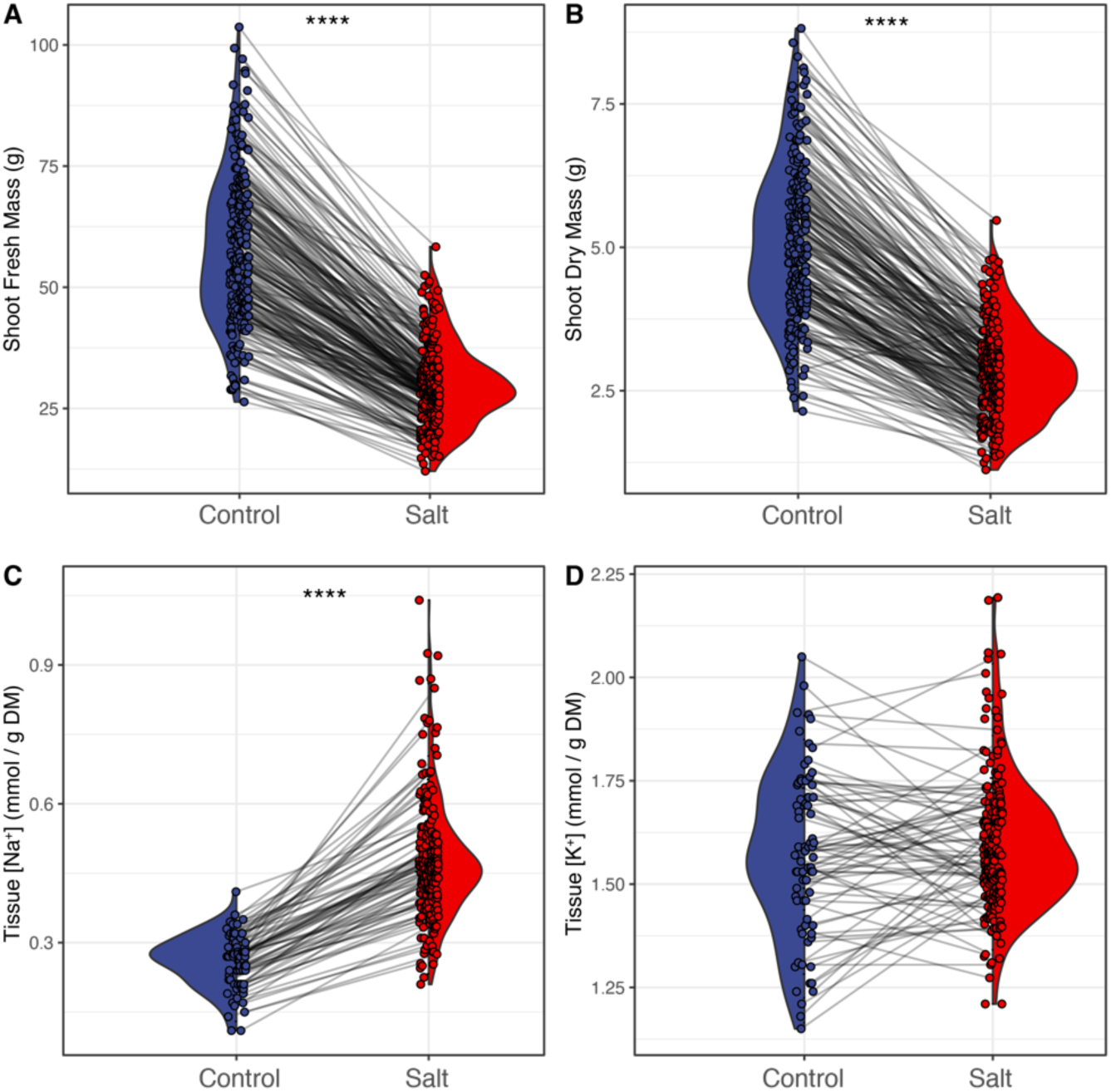
Salt stress reduced plant size while increasing sodium accumulation across *S. pimpinellifolium* accessions. Natural diversity panel of *S. pimpinellifolium* was germinated in the greenhouse conditions and exposed to an effective concentration of 0 or 200 mM NaCl at 21-24 days after sowing, corresponding to Control and Salt stress treatments respectively. After 2 weeks of salt stress treatment, the plants were harvested and evaluated for their **A)** fresh and **B)** dry mass. Additionally, leaves that developed during the salt stress treatment were collected and processed for **C)** sodium (Na^+^) and **D)** potassium (K^+^) contents. The individual lines in describe the change within a *S. pimpinellifolium* genotype observed between the treatments. The differences between treatments were tested using one-way ANOVA, and *, **, *** and **** indicate p-values below 0.05, 0.01, 0.001 and 0.0001 respectively.

To evaluate the relationships between the explored traits, we calculated the correlations between all measured traits (**Fig. S7**). Strong correlations were observed between shoot fresh mass (SFM), absolute growth rate, and transpiration rate across all intervals and treatments (r > 0.6, **Fig. S7**). Transpiration use efficiency was positively correlated with relative growth rate, but only under control conditions (0.3 < r < 0.6, **Fig. S7**). Interestingly, under salt stress conditions potassium and sodium accumulation were respectively negatively (-0.6 < r < -0.4) and positively (r ∼ 0.3) correlated with transpiration rate (**Fig. S7**). However, no consistent trends were observed when ion accumulation was overlaid over the relationship between shoot fresh mass under control and salt stress conditions (**Fig. S8**). To further explore how individual traits contribute to plant performance, we performed linear regression using individual traits to explain shoot fresh mass under control and salt stress conditions (**Fig. 4, Table S6**). Absolute growth rate, recorded in either of the three intervals, could explain beyond 60% variation in shoot fresh weight in both conditions. The second trait to explain most of the plant performance under control conditions was transpiration rate (75 to 64%). Interestingly, under salt stress conditions, the shoot fresh mass recorded under control conditions was the second trait to explain most (73%) of the plant performance. While ion accumulation explained only 20% and 5% of the variation in shoot fresh mass, it ranked among the traits that explained the least variance in overall plant performance (**Fig. 4, Table S6**). Together, these results suggest that the performance of *S.pimpinellifolium* shows wide variation in growth and response to salt stress. The plant performance under studied conditions cannot be easily explained by a single physiological process. Additionally, the observed traits show sufficient levels of variance and a broad sense heritability in the established population of *S. pimpinellifolium*, suggesting their high suitability for forward genetic studies.

**Figure 4.**
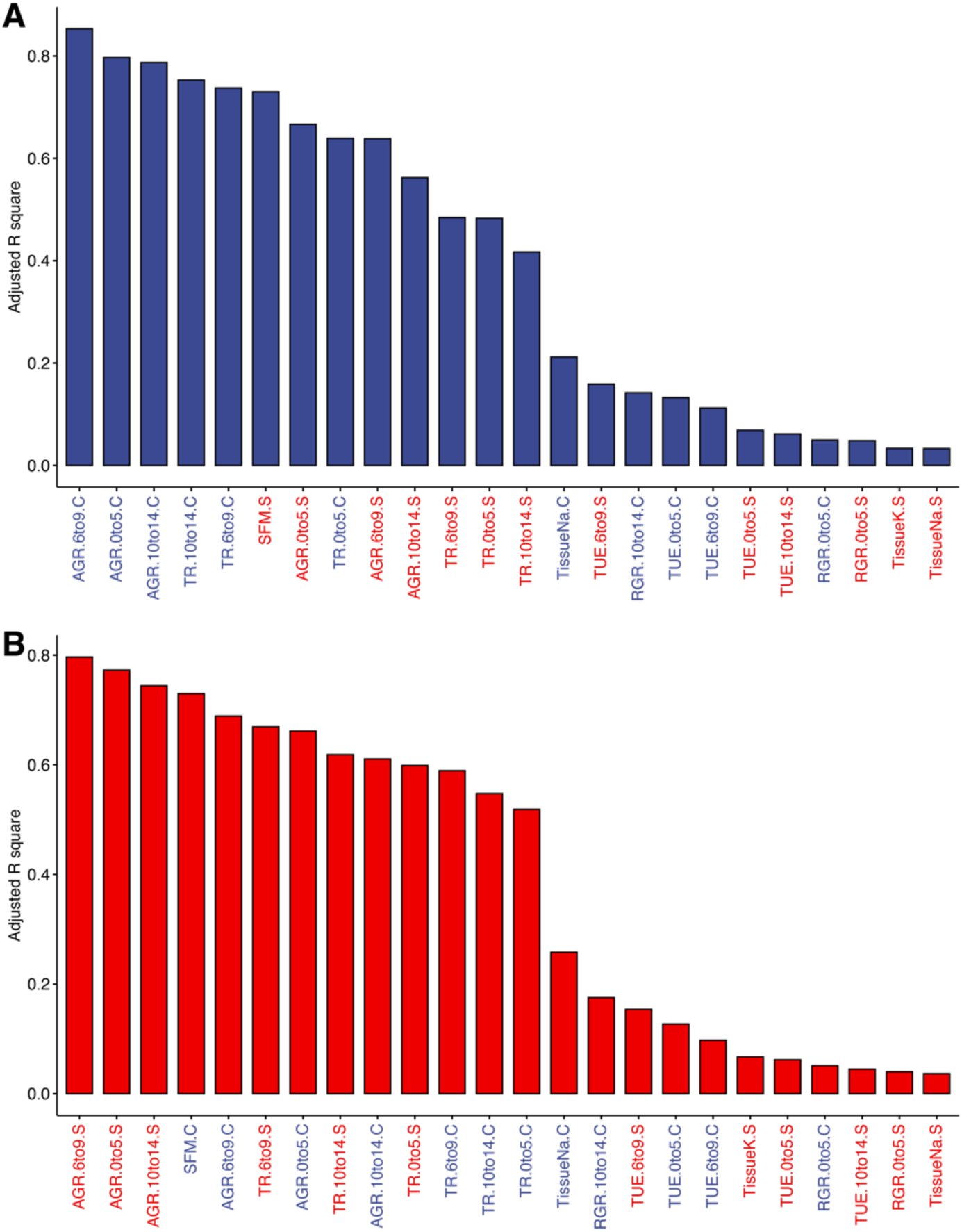
Plant performance under salt stress can be predicted from plant development and transpiration rate. All of the collected phenotypes over time of the greenhouse experiment were used to perform the linear regression modeling to explain shoot fresh mass (SFM) under **(A)** Control and **(B)** Salt stress conditions. The adjusted regression coefficient (R2) was examined for each measured trait, representing the fraction of explained variation inn SFM. The measured traits have been abbreviated as follows: AGR for Absolute Growth Rate; RGR for Relative Growth Rate, TR for Transpiration Rate, TUE for Transpiration Use Efficiency, TissueNa and TissueK for sodium and potassium ion accumulation in the shoot tissue respectively. The condition at which individual traits were measured is indicated with C or S for Control and Salt stress treatment respectively. The traits with not significant (p-value < 0.01) regression coefficients are not displayed

### Evaluating salinity tolerance of *S.pimpinellifolium* in the field

To evaluate agronomic performance under non-stress and salt stress conditions, 199 *S. pimpinellifolium* accessions and 1 *S. lycopersicum* (Heinz) were grown in the field conditions (**Fig. S9, Table S1**). The plants were exposed to mock treatment or salt stress at six weeks after germination, and imaged regularly using unmanned aerial vehicles (UAVs) (**Tables S7 - S9**) to estimate changes in the Projected Shoot Area (PSA), Modified Red Edge Normalized Difference Vegetation Index (MRENDVI), Photochemical Reflectance Index (PRI), and canopy temperature (**Fig. 5**). At 8 weeks after treatment imposition, all the field-grown plants were harvested and evaluated for shoot fresh mass, total fruit mass and number, as well as mature and immature fruit mass and number (**Fig. 6, Fig. S10**). The correlation between shoot fresh weight and projected shoot area for the field-grown plants was relatively high (r = 0.68 and 0.8 for control and salt stress-grown plants, respectively, **Fig. S10**), albeit lower than for the experiment performed under controlled conditions. This lower correlation could be stemming from the more advanced developmental stage of the imaged plants, and including plant architecture traits (height and volume), inferred from Canopy Height Model (Johansen et al., 2020) would improve this relationship.

**Figure 5.**
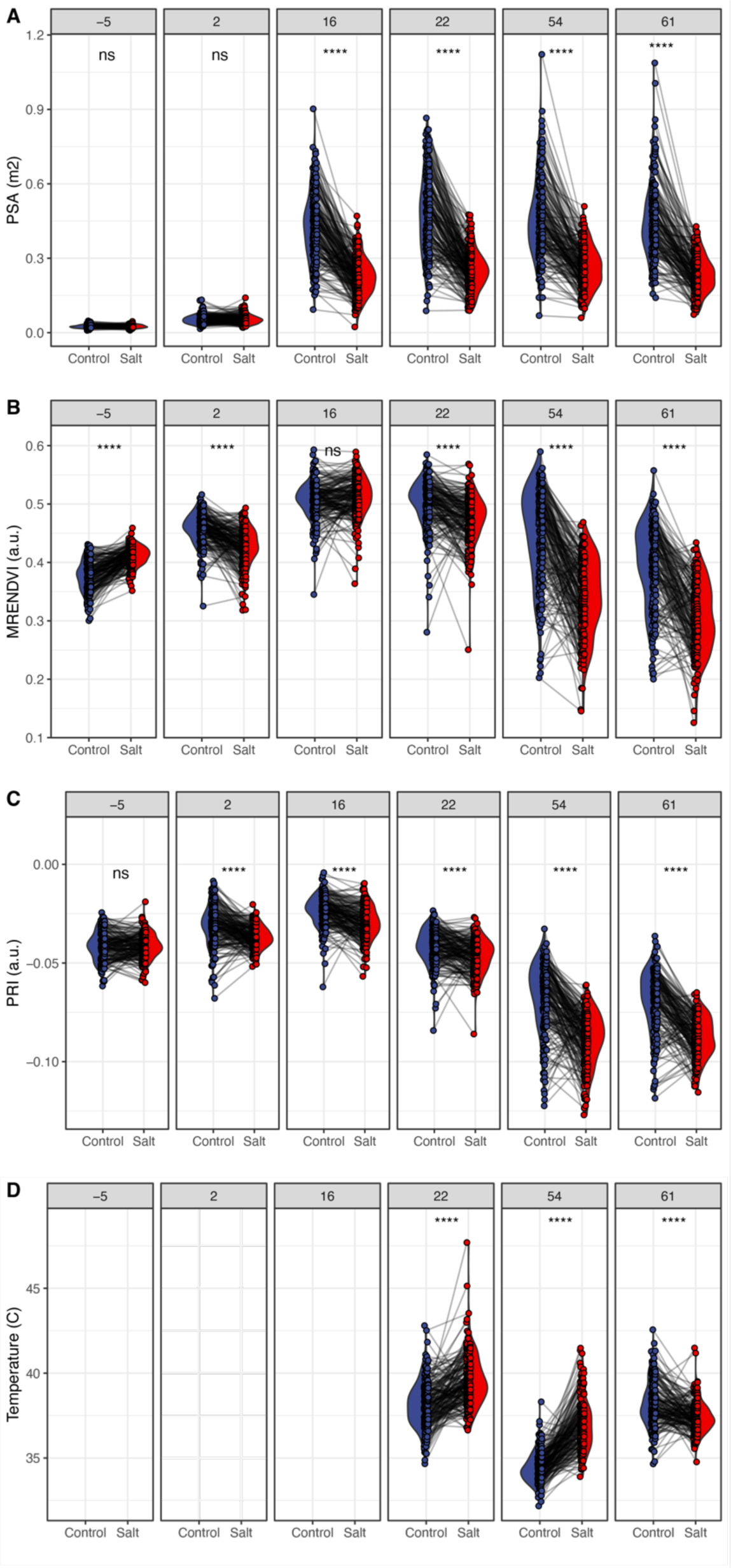
Salt reduced *S. pimpinellifolium* growth and yield, but not fruit maturation. Natural diversity panel of *S. pimpinellifolium* was germinated in the greenhouse conditions and exposed to an effective concentration of 0 or 200 mM NaCl at 21-24 days after sowing, corresponding to Control and Salt stress treatments respectively. The plants were monitored for 11 weeks during 6 UAV campaigns performed before and after treatment application for increase in the (**A)** projected shoot area, (**B)** MRENDVI, (**C)** PRI and **i**ndices and (**D)** canopy temperature. The numbers over individual graphs represent the days before or after salt stress application. The individual lines in describe the change within a genotype observed between the treatments. The differences between Control and Salt stress treatments were tested using ANOVA, and *, **, *** and **** indicate p-values below 0.05, 0.01, 0.001 and 0.0001 respectively.

**Figure 6.**
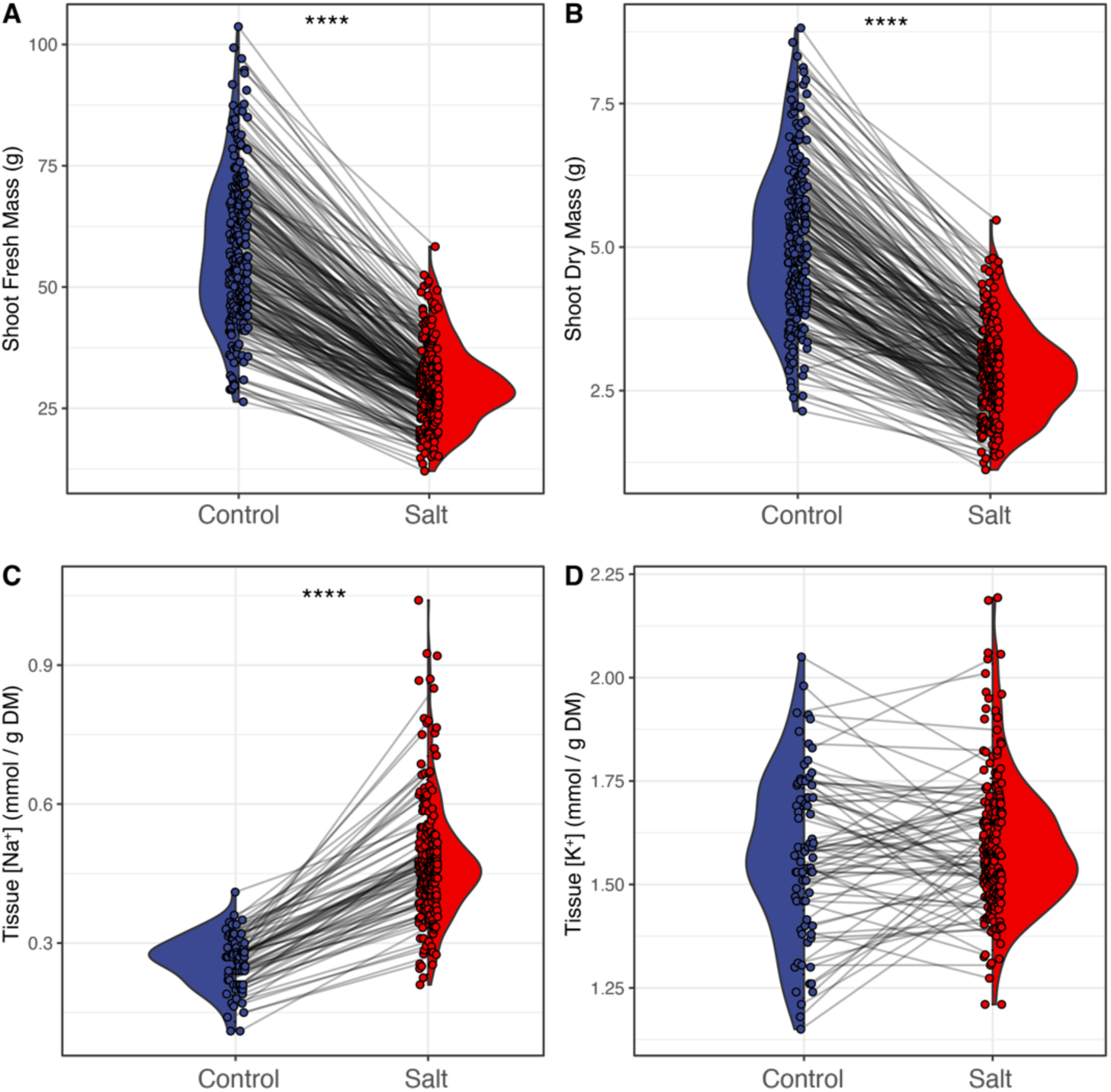
Salt treatment reduced plant productivity, but not fruit maturation. The *S. pimpinellifolium* plants exposed to control or salt stress treatment for over 60 days were manually harvested, and evaluated for (**A)** fresh mass and (**B)** total fruit yield. The collected fruits were further evaluated for maturity, and the data was used to calculate (**C)** genotype-specific fraction of mature fruit over the total fruit yield. Additionally, yield data was combined with shoot fresh mass data to calculate genotype-specific (**D)** harvest index (fruit yield / shoot fresh mass). The individual lines in describe the change within a genotype observed between the treatments. The differences between Control and Salt stress treatments were tested using ANOVA, and *, **, *** and **** indicate p-values below 0.05, 0.01, 0.001 and 0.0001 respectively.

A substantial increase in shoot area was observed over the first four imaging campaigns (5 days before to 22 days after stress imposition, **Fig. 5 A**), consistent with the exponential growth of plants during this period. Salt stress significantly decreased shoot area from 16 days after salt stress exposure onwards (**Fig. 5A**). The stagnation of projected shoot area from 22 days after stress imposition onwards likely reflects the limits of using plant area as a proxy for biomass. Including other architectural traits including leaf mass area, height, volume, could be considered to build a better biomass model, whereas including broader range of biochemical traits, such as chlorophyll or carotenoid content could serve as ancillary plant functional traits for better biomass predictions. The differences in MRENDVI between Control and Salt treated plants are not consistent prior to 16 days after salt stress imposition (**Fig. 5B**), after which salt-stressed plants exhibit consistently lower MRENDVI values compared to non-stressed plants. PRI differentiated between control and salt stressed plants immediately after salt stress imposition, with control plants consistently outperforming salt-stressed plants (**Fig. 5C**). Significant differences between treatments were also observed for canopy temperature for all available time points, with salt-stressed plants exhibiting higher temperatures (**Fig. 5D**). It is important to note that canopy temperature data does not account for differences in air temperature between individual campaign dates.

At the terminal harvest, salt stress negatively impacted all measured traits (**Tables S10 - S12, Fig. 6, Fig. S11**). The average shoot fresh mass was 67% lower in salt-stressed plants than in control plants. Salt-stressed plants have fewer immature and mature fruit (74.3% and 66.5% decrease in mean fruit number and mass, respectively), which translated to lower yields (79% and 74% decrease in immature and mature fruit yield, respectively). This is reflected in overall fruit numbers (71% decrease in fruit number) and yield (77% decrease in yield mass). The average fruit size also decreased in response to salt by 33% for both immature fruit (IFM) and mature fruit (MFM). Finally, the average harvest index (HI) decreased by 33% in salt-stressed plants relative to control plants.

Broad-sense heritability estimates were moderate for all the traits measured at the terminal harvest (*H^2^ =* 0.5 – 0.71, **Table S11**), except for shoot fresh mass, mature fruit number and yield under salt stress conditions (*H^2^ =* 0.24, 0.28, and 0.28, respectively). Heritability was lower under salt stress than control conditions, except for IFM, where heritability under salt stress (*H^2^ =* 0.57) was marginally higher than under control conditions (0.56) (**Table S11**). Heritability estimates for UAV vegetation indices (VI, i.e., PSA, MRENDVI and PRI) were relatively low (all < 0.5, except for MRENDVI under salt stress on 16 days after stress imposition), with many traits having broad-sense heritability values of 0. For canopy temperature, broad sense heritability values are close to 0 for all days under both conditions. These results indicate possible limited genetic influence on the variation in UAV derived indices within the experiment.

However, it is important to note that heritability is a complex concept, and it can be difficult to determine the extent to which genetic factors influence a particular trait. Therefore, we included all the measured traits and vegetation indices, including the ones with low heritability, in subsequent forward genetic analyses.

To examine the salt specific effects, seven stress tolerance indices were calculated for shoot fresh mass and total yield mass (**Fig.S12 - S13, Table S12**). Similarly to the trends observed for plants grown under greenhouse conditions, STI1, TOL, and SSI indices prioritized accessions for which performance under control and salt stress conditions are similar while MP, GMP, STI and SWP highlighted accessions with high performance under both conditions. Additionally, the indices calculated for yield data (**Fig.S13**) including STI1, SSI and SWP were prone to overestimating the effect of plants that performed particularly poorly under control conditions.

To examine the relationship between all measured traits, we calculated correlation coefficients (**Fig. S14**). Canopy temperature showed weak negative correlation (-0.3 < r < 0) with projected shoot area and MRENDVI, and high positive correlation (0.7 < r < 1) was observed between PRI and MRENDVI (**Fig. S14**). Terminally harvested traits, including shoot fresh mass and yield traits were also positively correlated with projected shoot area, MRENDVI and PRI (0 < r < 0.6, **Fig. S14**), and the correlations were strongest between the traits measured within the same treatment. To explore the individual trait contribution to overall plant performance, we executed linear regression for traits contributing to total yield (**Fig. 7**) and shoot fresh mass (**Fig. S15, Table S13**). The yield-related traits were amongst the highest ranked traits explaining total fruit yield under control and salt stress conditions, whereas shoot fresh mass explained 63 and 48% of the variation in yield for control and salt stressed plants respectively (**Fig. 7, Table S13**). The plant performance between the conditions was amongst the traits with medium contribution (20- 40%), whereas the indices (MRENDVI and PRI) explained the least of plant performance (< 20%) within and between the studied conditions (**Fig. 7, Table S13**). These results indicate that S. pimpinellifolium performance under both control and salt stress conditions are determined through plant development and the vegetative biomass, whereas the hyperspectral indices provide additional nuance to plant performance.

**Figure 7.**
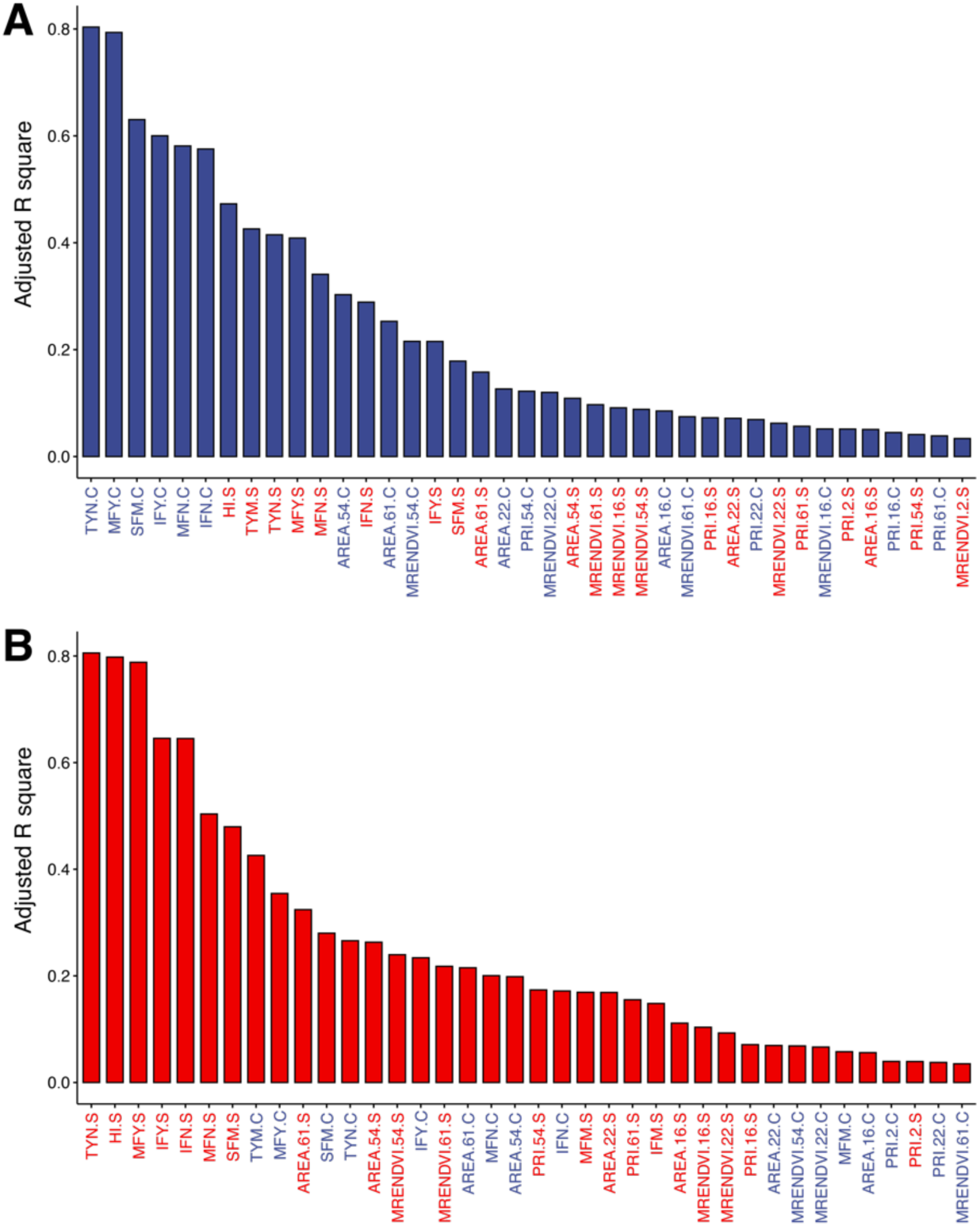
The plant yield is explained by plant size to a lesser degree under salt stress conditions. All of the collected phenotypes over time of the field experiment were used to perform the linear regression modeling to explain total yield mass (TYM) under **(A)** Control and **(B)** Salt stress conditions. The adjusted regression coefficient (R2) was examined for each measured trait, representing the fraction of explained variation in TYM. The measured traits have been abbreviated as follows: SFM for shoot fresh mass, AREA for projected shoot area, TYN for total yield number, MFY and IFY for mature and immature fruit yield, MFN and IFN for mature and immature fruit number respectively. The calculated Vegetation Indices were also included: PRI and MRENDVI that were calculated with multispectral data. The condition at which individual traits were measured is indicated with C or S for Control and Salt stress treatment respectively. The traits with not significant (p-value < 0.01) regression coefficients are not displayed

### Comparing salt stress performance between greenhouse and field experiments

To compare plant performance between greenhouse and field experiment, we examined the correlations between all the measured traits and VIs described within this study (**Fig. S15**). While the strongest correlations were observed between the traits recorded within the same experiment, we observed positive correlations between absolute growth rate, transpiration rate in greenhouse and projected shoot area in the field experiments. Under control conditions, the correlations were strongest for earlier time points, whereas under salt stress conditions, significant correlations were observed only for later time points (**Fig. S15**). The accumulation of sodium did not show consistent correlations with any of the performance traits under field conditions, whereas potassium accumulation under salt stress conditions was negatively correlated with plant performance under non-stress and salt stress conditions in the field (**Fig. S15**). To further evaluate the trait and VIs contribution to plant performance, we conducted a regression analysis for all studied traits and VIs to explain plant performance in greenhouse and field under non-stress and salt stress conditions (**Fig. S16, Table S14**). The traits recorded in a complementary experimental setup explained below 10% of the variance in shoot size or fruit yield (**Fig. S16, Table S14**). To identify specific genotypes that performed well under both greenhouse and field experiments, we further explored the correlations between shoot fresh mass across the experiments (**Fig. 8, Fig. S17 - S18**). While shoot fresh mass and yield were positively correlated under control and salt stress conditions across the experiment, no correlation was observed for STI1, SSI and SWP. We have identified LA1301, LA1374, LA2540, LA2836, and LA2974 as consistently high performing accessions across the experiments and conditions, whereas LA1279, LA1678, LA2340, LA3158 and LA3159 exhibited consistently poor performance. While the above results illustrate the challenges of comparing the performance of diversity panels between experimental setups, we were able to identify the genotypes with consistent performance across the experimental setups. Moreover, we have identified transpiration rate, absolute and relative growth rate and yield related traits to contribute to plant performance across complementary experimental setups. Identifying the genetic components underlying these traits will further accelerate development of climate resilient crops.

**Figure 8.**
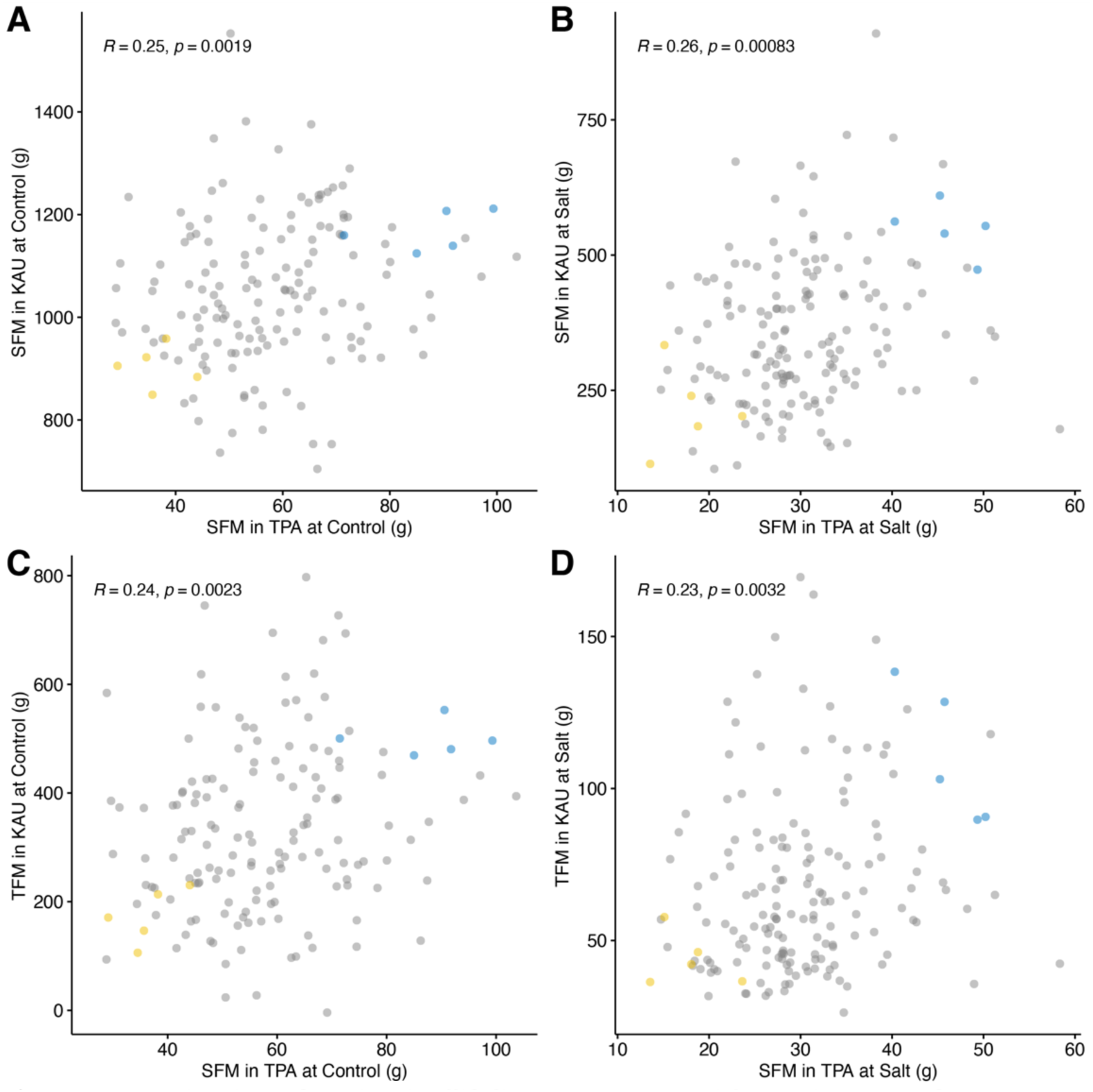
Examination of *S. pimpinellifolium* genotypes with consistent performance across experimental conditions. The performance of the plants across experimental conditions (greenhouse / TPA and field / KAU) was evaluated using the scatter plots and correlation analysis. The genotypes with best (blue points) and worst (yellow points) performance were identified based on their consistent performance between the experiments, based on (**A)** shoot fresh mass (SFM) under control and (**B)** salt stress conditions, as well as (**C)** total fruit mass (TFM) under control and (**D)** salt stress conditions. The pearson correlation coefficient and p-value is calculated for each scatter plot.

### Genome-Wide Association Study identifies new loci contributing to plant performance under salt stress conditions

To identify the genetic components underlying the variation in all observed traits, the collected phenotyping and genotypic data were used in Genome Wide Association Study (GWAS). The SNPs were further filtered to remove heterozygous markers and the ones represented by less than three genotypes (corresponding to MAF of 1.4 %). The GWAS was run including the kinship matrix (**Fig. S19**) similarly to the method described in (Awlia et al., 2021). As the accessions used in the greenhouse and field experiment differed slightly, we performed separate GWAS for each experiment.

Within the GWAS for traits recorded within the greenhouse experiment, we found a total of 345 associations that were above the Bonferroni threshold (> 7.15). The associations with shoot fresh and dry mass under salt stress conditions, sodium:potassium ratio were disregarded, based on the poor fit of the GWAS model as determined by QQ plots. After excluding these traits, the 345 associations were maintained above the Bonferroni threshold (**Table S15**). All the associations above the Bonferroni threshold were inspected for neighboring SNPs within 10 kbp, and grouped into 68 genetic loci. We subsequently evaluated each locus for the number of significantly associated SNPs, minor allele count and the number of traits associated with each locus, as this reduces the possibilities of the locus being a false-positive.

The three loci with most associated SNPs consisted of eight, seven and four significantly associated SNPs on chromosome 1 (**Table S16**), all associated with absolute growth under salt stress in the 1st and 3rd interval (0 to 5, and 10 to 14 days after stress imposition respectively) (**Fig. 9**). Additionally, SNPs with significance below the Bonferroni thresholds (-log10(*p-value*) > 5) were found in the same regions with the absolute growth rate under control conditions (**Fig. 9 B, D, F**). For Locus 1, the significantly associated SNPs were located in the coding regions of Spimp01g0026750 and Spimp01g0026760, both encoding homologue of ATP Binding Cassette (ABC) transporter C family member 14-like proteins in *S. lycopersicum* (**Table S16**). The linkage disequilibrium (LD) analysis revealed that SNPs from 77166348 to 77187538 were within LD of significantly associated SNPs, extending the region of putative candidate genes into Spimp01g0026770, encoding a gene with unknown function. For Locus 2, the significantly associated genes were located in Spimp01g0026870 and Spimp01g0026880, both encoding homologue of pathogen-inducible genes comprising the PR17 family homologue in *S. tuberosum* (**Table S16**), with Spimp01g0026860 and Spimp01g0026890 being located in LD, encoding F-box/kelch-repeat protein homologue and isoleucine--tRNA ligase respectively. The SNPs associated with Locus 3 were located between the gene coding regions of Spimp01g0029830 and Spimp01g0029840, encoding homologs of pheophytinase involved in chlorophyll breakdown during leaf senescence and zinc finger CCCH domain-containing protein 46, respectively. Other noteworthy associations with the greenhouse data included the locus identified for relative growth rate in the last interval of salt stress (10 to 14 days after stress imposition), with one SNP on chromosome 10 (**Fig. S20**), located within the coding regions of Spimp10g0288490 and Spimp10g0288500, both encoding homologue of universal stress protein (PHOS34). Additionally, we identified two loci significantly associated with sodium accumulation (**Fig. S21**), located on chromosomes 2 and 4 respectively, consisting of one and seven significantly associated SNPs, respectively. The SNP significantly associated with sodium accumulation on chromosome 2 was flanked by two uncharacterized genes (Spimp02g0051590 and Spimp02g0051600), whereas SNPs on chromosome 4 were positioned between Spimp04g0129310 and Spimp04g0129370, encoding uncharacterized genes (Spimp04g0129310, Spimp04g0129320, Spimp04g0129330, Spimp04g0129360 and Spimp04g0129370) and membralin-like proteins (Spimp04g0129340, Spimp04g0129350).

**Figure 9.**
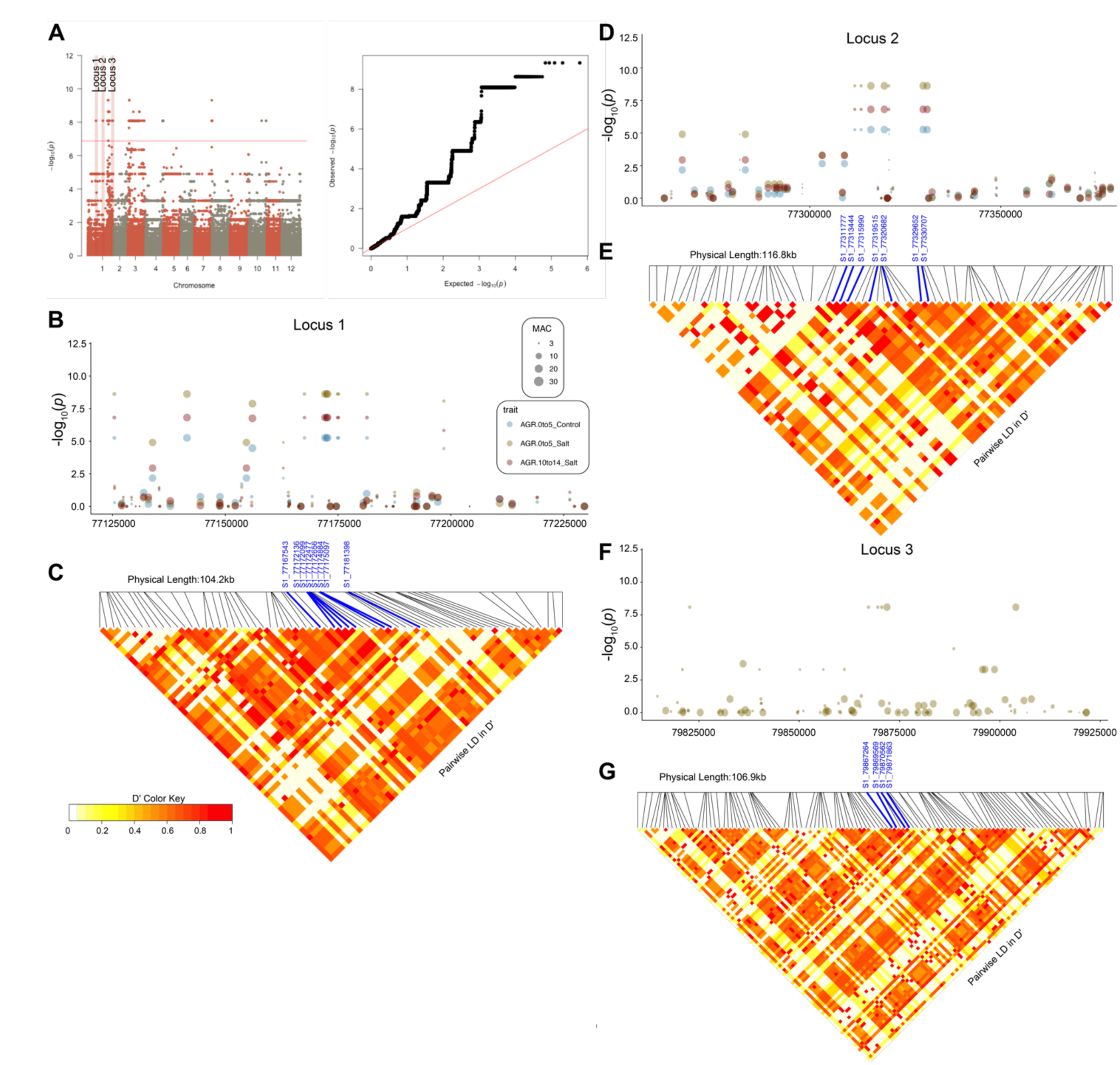
Natural variation in absolute growth rate during early exposure to salt stress in *S. pimpinellifolium* corresponds to three genetic loci. The Genome Wide Association Study was performed for 220 *S. pimpinellifolim* accessions using 708 545 SNPs. The ASReml GWAS model was applied on (A) absolute growth rate (AGR) in the 1st interval (0 to 5 days after treatment application). (B) The first identified locus consisted of eight SNPs (positions 77167543, 77172136, 177172099, 177172477, 177172656, 177174884, 77181398) on chromosome 1. The region within 100 kbp of identified SNPs was further inspected for associations with other traits. (C) The local linkage disequilibrium (LD) was calculated for all of the SNPs within the 100 kbp window. j(D) The second identified locus consisted of seven SNPs (positions 77311777, 77313444, 77315990, 77319515, 77320682, 77329652, 77330707) on chromosome 1. The region within 100 kbp of identified SNPs was further inspected for associations with other traits. (E) The local linkage disequilibrium (LD) was calculated for all of the SNPs within the 100 kbp window. (F) The third identified association consisted of four SNPs (positions 79871863, 79870562, 79869569, 79867264) on chromosome 1. The region within 100 kbp of identified SNPs was further inspected for associations with other traits. G) The local linkage disequilibrium (LD) was calculated for all of the SNPs within the 100 kbp window. The significantly associated SNP is highlighted in blue in LD heatmaps, whereas the LD between the individual SNPs is represented as a heatmap, with high intensity of red indicating strong linkage, and yellow hues indicating low linkage.

Within the GWAS for traits and VIs recorded in the field experiment, we found a total of 2,967 associations that were above the Bonferroni threshold (6.22). The associations with immature fruit yield under salt stress conditions, mature fruit mass under control conditions, mature fruit number under salt stress conditions and MRENDVI 2 days after treatment imposition under control conditions were disregarded, based on the poor fit of the GWAS model. No associations above Bonferroni threshold remained. Thus, we did not explore the identified associations associated with the field data.

## Discussion

Enhancing the sustainability of agriculture, particularly its dependency on freshwater, can be significantly advanced through investigations of stress tolerance mechanisms inherent in species closely related to crop plants (Bai and Lindhout, 2007; Zhang et al., 2017; Melino and Tester, 2023). The closest tomato relative, *S. pimpinellifolium (Bedinger et al., 2011)*, has previously been investigated for its potential for domestication (Zsögön et al., 2018), as well as plethora of abiotic and biotic factors (Ashrafi et al., 2009; Razali et al., 2018; Sullenberger et al., 2022). Recent assembly of a high-quality S. pimpinellifolium genome (Wang et al., 2020a) enables further genetic studies and development of natural diversity panels. Here, we present the establishment of a natural diversity panel consisting of 274 accessions of *S. pimpinellifolium* (**Fig. 1**). Our study has identified six accessions to be potentially misclassified as *S. pimpinellifolium* (LA0859, LA3158, LA3160, LA3161, LA3159 and LA2658), as they clustered together with cultivated tomatoes (**Fig. S2**). In total, we identified 20,325,817 SNPs within the *S. pimpinellifolium* diversity panel. Previous studies that explored natural diversity of *S.pimpinellifolium* identified fewer SNPs (3,125 SNPs in Celik et al., 2017, 7,414 SNPs in Blanca et al., 2012, 24,330 SNPs in Lin et al., 2019, 44,064 SNPs in Gibson and Moyle, 2020, 2,824,130 SNPs in Wang et al., 2020b), possibly due to aligning the reads to the *S. lycopersicum* genome rather than *S. pimpinellifolium*. Additionally, we observed population diversification across the North-South gradient, that is driving the diversity of the population (**Fig. 1**), similar to the trends observed previously (Gibson and Moyle, 2020).

The use of high-throughput phenotyping to study the responses to salt stress allowed us to characterize temporal changes within the studied population in response to stress (**Fig. 2**). Similar to other studies utilizing high-throughput phenotyping (Al-Tamimi et al., 2016; Awlia et al., 2021), most of the studied traits showed significant responses to salt stress (**Fig. 2**, **Fig. 3**, **Fig 5**, **Fig. 6**). The observed changes imposed by salt stress to PRI are consistent with previous study that reports PRI to be one of the most predictive vegetation index for retrieval of chlorophyll context (Angel and McCabe, 2022). Potassium accumulation (**Fig. 3D**) and the rate of fruit maturation (**Fig. S11**) did not show significant response to salt stress when considering the entire diversity panel, which is in line with previous observations (Martínez-Cuenca et al., 2020). Individual accessions that did show pronounced response in these specific traits could be identified (**Tables S5, S12**). While the experiment performed under greenhouse conditions was shorter, it provided more precise data, with an overall higher level of heritability, compared to the field experiment (**Tables S4, S11**). This was reflected by the significant candidate genes identified for the greenhouse experiment (**Table S16**), while no candidate genes could be identified for the field experiment using GWAS. Nevertheless, both experiments provided extensive insight into individual genotype performance under greenhouse and field conditions, which are arguably two most relevant conditions for the tomato agriculture (Heuvelink, 2005). Furthermore, the high-dimensional character of the collected data allowed us to explore the relationship within the phenotypic space, by using correlation analysis as well as linear regression models (**Fig. 4**, **Fig. 7, Fig. S7, Fig. S14, Tables S6, S13**). This analysis not only confirmed a strong relationship between all traits associated with plant growth, but also highlighted transpiration rate to explain plant size under control and salt stress conditions, whereas ion accumulation explained the least of the observed variation in plant performance (**Fig. 4**). Previous studies identified regional differences in vapor pressure as a strong selective factor within the *S. pimpinellifolium* population (Gibson and Moyle, 2020). While we did not identify any significant associations with transpiration rate (**Table S15**), the observed variation in transpiration rates can be further explored using individual accessions in more in-depth studies (**Table S5**).

Moreover, our results suggest a strong relationship between plant performance under stress and control conditions in both greenhouse and field experiments (**Fig. 4**, **Fig. 7, Fig. S7, Fig. S14**). This finding underscores the significance of plant vigor in determining plant performance under salt stress. The pivotal role of plant development in salt stress resilience has been echoed in previous studies (Julkowska et al., 2016; Awlia et al., 2021). Larger plants are supported by more extensive root systems, which allow water access over larger soil volume, which might help mitigate the effects of salt stress (Freschet et al., 2021). Moreover, larger shoot volume allows for the dilution of accumulated salt ions over larger biomass, reducing the relative ion accumulation (Reddy et al., 2017; van Zelm et al., 2020). As ion accumulation ranked among the traits explaining the lowest fraction of the variation in plant performance (**Fig. 4, Fig. S8**), we speculate that the developmental mechanisms that contribute to salt resilience act independently of ion accumulation.

By combining the variation in phenotype and genotype, we identified the putative genetic components underlying plant performance in the greenhouse conditions (**Fig. 9**). We have identified three high-confidence loci (**Table S16**), that were associated with absolute growth rate under control and salt stress conditions. The genes identified within the linkage disequilibrium region include ATP-binding cassette (ABC) transporter C-family (Spimp01g0026750 and Spimp01g0026760) and an unknown gene (Spimp01g0026770). ABC transporters have been previously shown to mediate cellular detoxification by facilitating the transport over the plasma or tonoplast membrane (Hwang et al., 2016). AtABCB14 has been reported to transport malate and affect stomatal conductance (Lee et al., 2008), whereas other ABC transporters have been reported in transport of plant hormones, such as abscisic acid (Kang et al., 2010), which plays a major role in salt stress responses. The transport properties of the putative ABC transporters associated with absolute growth rate of *S.pimpinellifolium* are yet to be characterized. The genes that were within locus 2 included F-box/kelch-repeat (FBK) protein (Spimp01g0026860) and pathogenesis related proteins (PR17, Spimp01g0026870, Spimp01g0026880). FBK proteins act as a part of SKP1-Cullin-F-box complex, marking proteins for degradation via the 26S proteasome (Abd-Hamid et al., 2020). As FBK proteins are directly binding to the substrate, they play a pivotal role in selective interactions with proteins targeted for degradation (Abd-Hamid et al., 2020). Over-expression of FBK proteins has been previously shown to both improve (Xu et al., 2014; Liu et al., 2020; Gao et al., 2022) and reduce abiotic stress tolerance (Yan et al., 2011). The expression of the identified FBK protein, its protein targets and effect on salt tolerance remain to be identified.

Genes that resided within the third locus associated with absolute growth rate included pheophytinase (Spimp01g0029830) and zinc finger CCCH domain-containing protein 46 (Spimp01g0029840)). Pheophytinase is an enzyme that facilitates an essential step in the chlorophyll catabolic pathway that leads to non-fluorescent degradation of chlorophyll, preventing accumulation of potentially reactive intermediates during leaf senescence (Schelbert et al., 2009). While no visible symptoms of senescence were observed within our experiment, pheophytinase might be playing an important role in photoprotective mechanisms during prolonged salt exposure, which can be revealed by screening the selected accessions of *S. pimpinellifolium* for their photosynthetic responses to salt stress, similar to the approach adopted in (Awlia et al., 2021). CCCH zinc-finger protein on the other hand is a member of a large family of proteins that consists of 80 members in *S. lycopersicum* (Xu, 2014). Many CCCH zinc-finger proteins contain nuclear localization sequence and DNA/RNA binding domains (Han et al., 2021), and regulate stress tolerance by direct activation of target genes, including SOS1 and ABA-responsive genes (Han et al., 2014; Bogamuwa and Jang, 2016; Seok et al., 2018). However, the interaction partners of Spimp01g0029840 and its effect on gene expression under salt stress conditions remain to be characterized in the future studies. None of the identified candidate genes overlapped with the loci identified in previous studies examining salt tolerance in tomato (Babu et al., 2012; Pan et al., 2012; Almeida et al., 2014; Csiszár et al., 2014; Bacha et al., 2015; Abouelsaad et al., 2016; Gharsallah et al., 2016; Jaime-Pérez et al., 2017; Bouzroud et al., 2018; Razali et al., 2018; Coyne et al., 2019; Bouzroud et al., 2020; Wang et al., 2020b; Mu et al., 2021; Romero-Aranda et al., 2021; Hong et al., 2023), which mainly focused on transcriptomic responses or ion accumulation in either wild or cultivated tomatoes. Our findings suggest that by using high-throughput phenotyping we can not only identify dynamic responses to stress, but also identify new components of stress resilience.

In summary, our study describes a development of a new, well genotyped natural diversity panel of a close crop relative, wild tomato, *S. pimpinellifolium.* We identified high and low performing accessions in greenhouse and field conditions. These specific genotypes can be used in future studies as allele donors for the further improvement of crop performance and bi-parental studies. Furthermore, our study demonstrates insights that can be gained from high throughput phenotyping studies, not only by using it as input for the GWAS, but also by exploring the relationship within the phenotype-space. The developed methods, data and identified genes of interest can further serve improvement of environmental resilience through identification of potential breeding targets and/or rootstocks for more sustainable agriculture.

## Acknowledgements

The authors would like to recognize TGRC for sharing the seeds of the initial germplasm that was used to develop the natural diversity panel described in this study, as well as KAUST greenhouse staff for their help in growing the germplasm during the propagation, and KAU team for their hard work to conduct the field experiments. Additionally, we would like to thank all of the members from The Salt Lab who helped with the manual harvesting during the fieldwork. Financial support from KAUST to Prof. Mark Tester is gratefully acknowledged. The Australian Plant Phenomics Facility is supported by the Australian Government under the National Collaborative Research Infrastructure Strategy (NCRIS).

## Supplementary material

**Figure S1. The density of SNP identified in S. pimpinellifolium diversity panel distributed over 12 chromosomes.** All the mapped SNPs (including heterozygous SNPs) are plotted over their respective position within the *S. pimpinellifolium* calculated in a bin size of 100 kb. The yellow and blue shades represent high and low number of identified SNPs respectively.

**Figure S2. Population structure of wild and cultivated tomatoes.** Population structure (from *K* = 2 to *K* = 10) of 482 *S. pimpinellifolium*, *S. lycopersicum* and *S. lycopersicum var. cerasiforme* samples estimated with sNMF. Each bar represents a sample and the bars are filled by colors representing the likelihood of membership to each ancestry.

**Figure S3. Design of the greenhouse in The Plant Accelerator, for controlled environment phenotyping. A)** Model of the two TPA® smarthouses and phenotyping stations used in this experiment. WWS: weighing and watering station, Thermal IR: thermal infrared imaging station*, RGB: Red-Blue-Green imaging station, NIR: near infrared imaging*; F: fluorescence imaging*. (*) Imaging stations were not functional for this experiment. Image credit: Heno Hwang. **B)** *S. pimpinellifolium* plants growing in the NW smarthouse. **C)** Plot of the design showing the allocation of accessions to main plots, each main plot being comprised of two consecutive carts. The two salt treatment were randomized to the two consecutive carts within a main plot. The accessions are numbered from 1 to 220. The cells coloured orange are a complete replicate the 220 accessions within a greenhouse, and the cells coloured blue are the second replicates of the 68 accessions that had two replicates within a greenhouse. The white cells are the evaporation pots.

**Figure S4. The relationship between shoot fresh mass and projected shoot area in *S. pimpinellifolium* plants grown under environmentally controlled conditions.** The 220 *S. pimpinellifolium* accessions were grown under greenhouse conditions with and without exposure to salt stress for 2 weeks. The shoot fresh weight, recorded at the last day of the experiment, was compared to projected shoot area, recorded at the same day. The Spearman correlation was calculated for each treatment, together with the correlation p-value. The individual points represent genotype mean at individual condition. The lines represent the linear regression between the two traits.

**Figure S5. Data complexity was reduced by summarizing plant growth and transpiration rates into three intervals.** The change in the trait value observed over the course of two weeks of Control or Salt stress treatment (Figure 2), was summarized by calculating a genotypic mean values for three intervals: 0 to 5 days, 6 to 9 days, and 10 to 14 days. We examined stress-induced changes in **A)** Absolute Growth Rate (AGR), **B)** Relative Growth Rate (RGR), **C)** Transpiration Rate and **D)** Transpiration Use Efficiency (TUE). The individual lines in describe the change within a genotype observed between the treatments. The differences between treatments were tested using one-way ANOVA, and *, **, *** and **** indicate p-values below 0.05, 0.01, 0.001 and 0.0001 respectively.

**Figure S6. Evaluation of the *S.pimpinellifolium* accessions and identification of the most tolerant genotypes based on shoot fresh weight of plants grown under controlled environmental conditions.** The performance of *S. pimpinellifolium* accessions was evaluated based on the Shoot Fresh Mass (SFM) accumulated at the end of experiment (2 weeks after initial Control / Salt stress treatment). **A)** The relation between SFM of each genotype under Control and Salt stress conditions was examined using scatter plot, and the Spearman correlation coefficient (R) was calculated, along with the correlation p-value. Subsequently, we highlighted the accessions that showed highest and lowest performance based on the 7 stress tolerance indices: **B)** Salt tolerance index (STI = Salt / Control), **C)** Tolerance index (TOL = Control - Salt), **D)** Mean productivity index (MP = (Control + Salt) / 2), **E)** Geometric Mean Productivity (GMP = square root of (Control x Salt), **F)** Stress Susceptibility Index (SSI = ((Control - Salt) / Control)/ ((population mean Control - population mean Satl) / population mean Control), **G)** Stress Tolerance Index (STI2 = (Control x Salt) / population mean Control^2^), **H)** Stress Weighted Performance Index (SWP = Salt / square root of Control). The color scale for each index is contained within each graph, with red and blue points representing the best and poorest performing genotypes, respectively.

**Figure S7. Correlations between the traits recorded in the greenhouse experiment.** The pearson’s correlation coefficients were calculated between each combination of traits. Positive and negative correlations are depicted using orange and blue hues, respectively. Non-significant correlations (p-value > 0.05) are indicated with an X. The individual traits are abbreviated with AGR for absolute growth rate, RGR for relative growth rate, TR for transpiration rate, TUE for transpiration use efficiency, TissueK and TissueN for potassium and sodium accumulation, and SFM for shoot fresh mass. Traits recorded under control and salt stress conditions are indicated with C or S, respectively, at the end of each trait.

**Figure S8. Evaluation of the *S.pimpinellifolium* accessions for sodium and potassium accumulation.** The performance of 220 *S. pimpinellifolium* accessions was evaluated based on the shoot fresh mass (SFM) accumulated at the end of experiment (2 weeks after initial Control / Salt stress treatment). **A)** The relation between SFM of each genotype under Control and Salt stress conditions was examined using scatter plot, and the A) leaf sodium (Na^+^) and B) potassium (K^+^) accumulation is superimposed on the plant performance using the color scale. Color scale for each element is included within each plot, with red and blue points representing high and low ion accumulation respectively.

**Figure S9. Design of the field experiment. A)** Layout of field site **B)** Aerial view of the field site, with different plots labelled and due north indicated.

**Figure S10. The relationship between shoot fresh mass and projected shoot area in *S. pimpinellifolium* plants grown under field conditions.** The 119 *S. pimpinellifolium* accessions were grown under field conditions with and without exposure to salt stress for 11 weeks. The shoot fresh weight, recorded at the last day of the experiment, was compared to projected shoot area, recorded at the same day. The Spearman correlation was calculated for each treatment, together with the correlation p-value. The individual points represent genotype mean at individual condition. The lines represent the linear regression between the two traits.

**Figure S11. Salt treatment reduced plant productivity, but not fruit maturation.** The *S. pimpinellifolium* plants exposed to control or salt stress treatment for over 60 days were manually harvested, and evaluated for **A)** total fruit number (TFN) **B)** mature fruit yield (g), **C)** Mature fruit number (MFN), **D)** Mature fruit mass (g), **E)** Immature Fruit Yield (g), **F)** Immature fruit number (IFN), **G)** Immature fruit mass (g) and **H)** ratio of mature to immature fruit yield (MFY / IFY). The individual lines in describe the change within a genotype observed between the treatments. The differences between Control and Salt stress treatments were tested using ANOVA, and *, **, *** and **** indicate p-values below 0.05, 0.01, 0.001 and 0.0001 respectively.

**Figure S12. Evaluation of the *S.pimpinellifolium* accessions and identification of the most tolerant genotypes based on shoot fresh weight of plants grown under field conditions.** The performance of 199 *S. pimpinellifolium* accessions was evaluated based on the shoot fresh mass (SFM) accumulated at the end of experiment (11 weeks after initial Control / Salt stress treatment). **A)** The relation between SFM of each genotype under Control and Salt stress conditions was examined using scatter plot, and the Spearman correlation coefficient (R) was calculated, along with the correlation p-value. Subsequently, we highlighted the accessions that showed highest and lowest performance based on the 7 stress tolerance indices: **B)** Salt tolerance index (STI = Salt / Control), **C)** Tolerance index (TOL = Control - Salt), D) Mean productivity index (MP = (Control + Salt) / 2), **E)** Geometric mean productivity (GMP = square root of (Control x Salt), **F)** Stress Susceptibility Index (SSI = ((Control - Salt) / Control)/ ((population mean Control - population mean Satl) / population mean Control), **G)** Stress Tolerance Index (STI2 = (Control x Salt) / population mean Control^2^), **H)** Stress Weighted Performance Index (SWP = Salt / square root of Control). The color scale for each index is contained within each graph, with red and blue points representing the best and poorest performing genotypes.

**Figure S13. Evaluation of the *S.pimpinellifolium* accessions and identification of the most tolerant genotypes based on total yield mass of plants grown under field conditions.** The performance of 199 *S. pimpinellifolium* accessions was evaluated based on the total yield mass (TYM) accumulated at the end of experiment (11 weeks after initial Control / Salt stress treatment). **A)** The relation between TYM of each genotype under Control and Salt stress conditions was examined using scatter plot, and the Spearman correlation coefficient (R) was calculated, along with the correlation p-value. Subsequently, we highlighted the accessions that showed highest and lowest performance based on the 7 stress tolerance indices: **B)** Salt tolerance index (STI = Salt / Control), **C)** Tolerance index (TOL = Control - Salt), D) Mean productivity index (MP = (Control + Salt) / 2), **E)** Geometric mean productivity (GMP = square root of (Control x Salt), **F)** Stress Susceptibility Index (SSI = ((Control - Salt) / Control)/ ((population mean Control - population mean Satl) / population mean Control), **G)** Stress Tolerance Index (STI2 = (Control x Salt) / population mean Control^2^), **H)** Stress Weighted Performance Index (SWP = Salt / square root of Control). The color scale for each index is contained within each graph, with red and blue points representing the best and poorest performing genotypes.

**Figure S14. Correlations between the traits recorded in the field experiment.** The pearson’s correlation coefficients were calculated between each combination of traits. Positive and negative correlations are depicted using orange and blue hues, respectively. Non-significant correlations (p-value > 0.05) are indicated with an X. The individual traits are abbreviated with AREA for projected shoot area, MRENDVI for Modification Red Edge Normalised Difference Vegetation Index, PRI for Photochemical Reflectance Index, Temp. for canopy temperature, SFM for shoot fresh mass, MFY for mature fruit yield (g), MFN for mature fruit number, MFM for average mature fruit mass (g), IFY for immature fruit yield (g), IFN for immature fruit number, IFM for average immature fruit mass (g), TYM for total yield mass (g) and TYN for total fruit number. Traits recorded at various time points are indicated with a number following the trait description, corresponding to the days after salt stress imposition. Traits recorded under control and salt stress conditions are indicated with C or S, respectively, at the end of each trait.

**Figure S15. Correlations between the traits recorded in the greenhouse and field experiment.** The pearson’s correlation coefficients were calculated between each combination of traits. Positive and negative correlations are depicted using orange and blue hues, respectively. Non-significant correlations (p-value > 0.05) are indicated with an X. The individual traits are abbreviated with AGR for absolute growth rate, RGR for relative growth rate, TR for transpiration rate, TUE for transpiration use efficiency, TissueK and TissueN for potassium and sodium accumulation, and SFM for shoot fresh mass, AREA for projected shoot area, MRENDVI for Modification Red Edge Normalised Difference Vegetation Index, PRI for Photochemical Reflectance Index, Temp. for canopy temperature, SFM for shoot fresh mass, MFY for mature fruit yield (g), MFN for mature fruit number, MFM for average mature fruit mass (g), IFY for immature fruit yield (g), IFN for immature fruit number, IFM for average immature fruit mass (g), TYM for total yield mass (g) and TYN for total fruit number. Traits recorded at various time points are indicated with a number following the trait description, corresponding to the days after salt stress imposition. Traits recorded under greenhouse conditions carry prefix TPA (for The Plant Accelerator), whereas traits recorded under field conditions carry prefix KAU (for King Abdulaziz University farm). Traits recorded under control and salt stress conditions are indicated with C or S, respectively, at the end of each trait.

**Figure S16. The regression analysis of plant performance under greenhouse and field conditions.** All of the collected phenotypes over time of the greenhouse (TPA) and field (KAU) experiment were used to perform the linear regression modeling to explain **A)** Shoot fresh mass under control and **B)** salt stress conditions recorded under greenhouse conditions; **C)** shoot fresh mass under control and **D)** salt stress conditions and **E)** total yield mass under control and **F)** salt stress conditions under field conditions. The adjusted regression coefficient (R2) was examined for each measured trait, representing the fraction of explained variation in plant performance. The individual traits are abbreviated with AGR for absolute growth rate, RGR for relative growth rate, TR for transpiration rate, TUE for transpiration use efficiency, TissueK and TissueN for potassium and sodium accumulation, and SFM for shoot fresh mass, AREA for projected shoot area, MRENDVI for Modification Red Edge Normalised Difference Vegetation Index, PRI for Photochemical Reflectance Index, Temp. for canopy temperature, SFM for shoot fresh mass, MFY for mature fruit yield (g), MFN for mature fruit number, MFM for average mature fruit mass (g), IFY for immature fruit yield (g), IFN for immature fruit number, IFM for average immature fruit mass (g), TYM for total yield mass (g) and TYN for total fruit number. Traits recorded under greenhouse conditions carry prefix TPA (for The Plant Accelerator), whereas traits recorded under field conditions carry prefix KAU (for King Abdulaziz University farm). Traits recorded under control and salt stress conditions are indicated with C or S, respectively, at the end of each trait. The traits with not significant (p-value < 0.01) regression coefficients are not displayed.

**Figure S17. Evaluation of the *S.pimpinellifolium* accessions and identification of the most tolerant genotypes based on shoot fresh mass of plants grown under controlled environment and field conditions.** The performance of 199 *S. pimpinellifolium* accessions that were included in both controlled environment and field conditions screening was evaluated based on the shoot fresh mass (SFM) accumulated at the end of experiment (2 weeks for controlled conditions, 11 weeks for field experiment). The relation between SFM of each genotype under field and controlled environment experiment at The Plant Accelerator (TPA) was evaluated under **A)** Control and **B)** Salt stress conditions. Additionally, we evaluated the similarities in genotype performance using the various stress indices: **C)** Salt tolerance index (STI = Salt / Control), **D)** Tolerance index (TOL = Control - Salt), **E)** Mean productivity index (MP = (Control + Salt) / 2), **F)** Geometric mean productivity (GMP = square root of (Control x Salt), **G)** Stress Susceptibility Index (SSI = ((Control - Salt) / Control)/ ((population mean Control - population mean Satl) / population mean Control), **H)** Stress Tolerance Index (STI2 = (Control x Salt) / population mean Control^2^), **I)** Stress Weighted Performance Index (SWP = Salt / square root of Control). Each relationship between control and field conditions was examined using scatter plot, and the Spearman correlation coefficient (R) was calculated, along with the correlation p-value.

**Figure S18. Evaluation of the *S.pimpinellifolium* accessions and identification of the most tolerant genotypes based on shoot fresh mass of plants grown under controlled environment and total yield mass of plants grown under field conditions.** The performance of 199 *S. pimpinellifolium* accessions that were included in both controlled environment and field conditions screening was evaluated based on the shoot fresh mass (SFM) for controlled environmental conditions and total fruit mass (TFM) for field conditions accumulated at the end of experiment (2 weeks for controlled conditions, 11 weeks for field experiment). The relation between TFM and SFM of each genotype under field and controlled environment experiment at The Plant Accelerator (TPA) was evaluated under **A)** Control and **B)** Salt stress conditions. Additionally, we evaluated the similarities in genotype performance using the various stress indices: **C)** Salt tolerance index (STI = Salt / Control), **D)** Tolerance index (TOL = Control - Salt), **E)** Mean productivity index (MP = (Control + Salt) / 2), **F)** Geometric mean productivity (GMP = square root of (Control x Salt), **G)** Stress Susceptibility Index (SSI = ((Control - Salt) / Control)/ ((population mean Control - population mean Satl) / population mean Control), **H)** Stress Tolerance Index (STI2 = (Control x Salt) / population mean Control^2^), **I)** Stress Weighted Performance Index (SWP = Salt / square root of Control). Each relationship between control and field conditions was examined using scatter plot, and the Spearman correlation coefficient (R) was calculated, along with the correlation p-value.

**Figure S19. The kinship matrix for *S. pimpinellifolium* accessions used as a co-factor for Genome Wide Association Study.** The kinship matrix was calculated in GAPIT using 708,545 SNPs that underwent filtering and were subsequently used in GWAS. The kinship between individual accessions is represented as a Z-score value, with red and white hues representing high and low level of kinship. The dendrogram represents hierarchical clustering of 199 genotypes of *S. pimpinellifolium* that were successfully sequenced and used in phenotyping experiments.

**Figure S20. Natural variation in relative growth rate and salt tolerance index under salt stress in *S. pimpinellifolium* corresponds to one genetic locus.** The Genome Wide Association Study was performed for 220 *S. pimpinellifolim* accessions using 708 545 SNPs. The ASReml GWAS model was applied on **A)** relative growth rate (RGR) in the 3rd interval, and **B)** Salt Tolerance Index (STI = S / C) calculated using RGR for 3rd interval (10 to 14 days after stress application). The identified association consisted of one SNP (position 1198094) on chromosome 9. **C)** The associations in both traits were inspected for other SNPs within the 100 kbp window. **D)** The local linkage disequilibrium (LD) was calculated for all of the SNPs within the 100 kbp window. The significantly associated SNP is highlighted in blue in LD heatmaps, whereas the LD between the individual SNPs is represented as a heatmap, with high intensity of red indicating strong linkage, and yellow hues indicating low linkage.

**Figure S21. Natural variation in sodium accumulation in *S. pimpinellifolium* corresponds to genetic variation distributed over two loci.** The Genome Wide Association Study was performed for 220 *S. pimpinellifolim* accessions using 708 545 SNPs. The ASReml GWAS model was applied on sodium (Na^+^) accumulation in the leaf tissue recorded under **A)** Control and **B)** Salt stress conditions. **C)** The first locus was identified on chromosome 2, position 9912134. The association was inspected for other SNPs within the 100 kbp window. **D)** The local linkage disequilibrium (LD) was calculated for all of the SNPs within the 100 kbp window. **E)** The second locus was identified on chromosome 4, and consisted of seven significantly associated SNPs (positions 18514133, 18509635, 18498875, 18495923, 18488327, 18487801 and 18486112). The region was further inspected for associations within 100 kbp window. F) The local LD was calculated for all of the SNPs within the 100 kbp window. The significantly associated SNPs are highlighted in blue in LD heatmaps, whereas the LD between the individual SNPs is represented as a heatmap, with high intensity of red indicating strong linkage, and yellow hues indicating low linkage.

**Supplemental Table S1. S.pimpinellifolium accessions used for Illumina sequencing and phenotyping experiments.** The sequencing depth is reported for each of the accession as a coverage on a basis of an estimated genome size of 0.9G. The accessions’ inclusion in The Plant Accelerator experiment (greenhouse conditions) and field experiment (Hadda Al Sham) is indicated with 1, whereas when the accession was omitted for a specific experiment, it is indicated with 0.

**Supplemental Table S2. Number of SNPs retained after each filtering step.**

**Supplemental Table S3. Population structure of S. pimpinellifolium samples.** Each accession is listed with their respective likelihood of membership to each ancestry (q1 - q5). The maximal contribution to specific cluster is listed in the last column.

**Supplemental Table S4. Overview of the effect of salt stress on mean trait value and broad-sense heritability in The Plant Accelerator experiment.** The traits recorded non-destructively are presented for each interval (0 to 5, 6 to 9 and 10 to 14 days after salt stress application). The results from a one-way ANOVA are listed for each trait to determine the population-wide difference between control and salt stress treated plants. Traits recorded at the terminal harvest (Shoot fresh mass, Leaf Na+ and K+) were analyzed only for the significant difference between control and salt stress treated plants (ANOVA p-value). The effect of salt stress was estimated based on the difference between the mean under control and salt stress. Broad-sense heritability (H2) was calculated for each interval, treatment and trait.

**Supplemental Table S5. TPA data and Indices. The collected data over the entire Plant Accelerator experiment.** The data collected over multiple time points is summarized into three intervals (0-5, 6-9 and 10-14 days after stress application), recorded under control (C) or salt stress (S) conditions. Based on the projected shoot area, Absolute and Relative Growth Rate were calculated (AGR and RGR respectively). Transpiration Use Efficiency (TUE) was calculated by using the change in the projected shoot area and transpiration for each interval (see formula in Materials & Methods section). At the last day of experiment, Shoot Fresh Mass (SFM) was recorded, and sodium (Na) and potassium (K) accumulations were measure. The stress indices were calculated using shoot fresh mass. In total - seven individual indices were calculated: Salt tolerance index (STI1 = S/C); Tolerance index (TOL = C-S); Mean productivity index (MP = (C+S)/2); Geometric mean productivity (GMP = sqrt(C*S)); Stress susceptibility index (SSI = ((C-S)/C) / ((C.mean-S.mean)/C.mean)); Stress tolerance index (STI2 = C*S/C.mean); Stress weighted performance index (SWP = S / sqrt(C)).

**Supplemental Table S6. Regression analysis between all traits recorded in the greenhouse-based experiment.** The first tab in this table contains the regression coefficients (R2) between each combination of traits, while the second contains adjusted R2. The third tab contains the p-values for each trait combination, whereas the fourth tab contains the residual standard error (RSE). The regression coefficients for which pvalue was > 0.01 were removed from both Rsquare and Adj.Rsquare tables. The measured traits have been abbreviated as follows: AGR for Absolute Growth Rate; RGR for Relative Growth Rate, TR for Transpiration Rate, TUE for Transpiration Use Efficiency, TissueNa and TissueK for sodium and potassium ion accumulation in the shoot tissue respectively. The condition at which individual traits were measured is indicated with C or S for Control and Salt stress treatment, respectively.

**Supplemental Table S7. Description of narrowband vegetation indices estimated from hyperspectral imagery.**

**Supplemental Table S8. Schedule of ground truth measurements and phenotyping campaigns using different cameras for the field site experiment.** The green-shaded cells correspond to the times at which ground-truthing was performed for this particular trait. The dates are in DD-MM-YR format.

**Supplemental Table S9. Description of UAV-based measurements, the cameras used to achieve them and the ground-truthing method employed to calibrate/validate them.**

**Supplemental Table S10. Summary of phenotypic traits determined at harvest.**

**Supplemental Table S11. Overview of the effect of salt stress on mean trait value and broad-sense heritability in the field experiment.** The traits recorded using UAVs are presented for each day after salting (DAS) for plant area, MRENDVI, PRI and canopy temperature under control and salt stressed on different days. The results from a two-way ANOVA followed by Tukey’s highly significant difference grouping (p < 0.05), are represented with letters designating groups that are significantly different. Traits recorded at the terminal harvest (TH) were anayzed only for the significant difference between control and salt stress treated plants (ANOVA p-value). The effect of salt stress was estimated based on the difference between the mean under control and salt stress. Broad-sense heritability (H2) was calculated for each day, treatment and trait.

**Supplemental Table S12. Field data and Indices.** The collected over the entire field experiment. The data collected using the UAVs over multiple timepoints is indicated using days before or after salting (DBS and DAS, respectively). The data used to calculate all the stress indices for shoot fresh mass (SFM), total yield mass (TYM) and harvest index (HI). For each trait - seven individual indices were calculated: Salt tolerance index (STI1 = S/C); Tolerance index (TOL = C-S); Mean productivity index (MP = (C+S)/2); Geometric mean productivity (GMP = sqrt(C*S)); Stress susceptibility indes (SSI = ((C-S)/C) / ((C.mean-S.mean)/C.mean)); Stress tolerance index (STI2 = C*S/C.mean); Stress weighted performance index (SWP = S / sqrt(C)).

**Supplemental Table S13. Regression analysis between all traits recorded under field conditions.** The first tab in this table contains the regression coefficients (R2) between each comination of traits, while the second contains adjusted R2. The third tab contains the p-values for each trait combination, whereas the fourth tab contains the residual standard error (RSE). The regression coefficients for which pvalue was > 0.01 were removed from both Rsquare and Adj.Rsquare tables. The individual traits are abbreviated with AREA for projected shoot area, MRENDVI for Modification Red Edge Normalized Difference Vegetation Index, PRI for Photochemical Reflectance Index, Temp. for canopy temperature, SFM for shoot fresh mass, MFY for mature fruit yield (g), MFN for mature fruit number, MFM for average mature fruit mass (g), IFY for immature fruit yield (g), IFN for immature fruit number, IFM for average immature fruit mass (g), TYM for total yield mass (g) and TYN for total fruit number. Traits recorded at various time points are indicated with a number following the trait description, corresponding to the days after salt stress imposition. Traits recorded under control and salt stress conditions are indicated with C or S, respectively, at the end of each trait.

**Supplemental Table S14. Regression analysis between all traits recorded across the experiments.** The first tab in this table contains the regression coefficients (R2) between each combination of traits, while the second contains adjusted R2. Third tab contains the p-values for each trait combination, whereas fourth tab contains the residual standard error (RSE). The regression coefficients for which pvalue was > 0.01 were removed from both Rsquare and Adj.Rsquare tables. The measured traits have been abbreviated as follows: AGR for Absolute Growth Rate; RGR for Relative Growth Rate, TR for Transpiration Rate, TUE for Transpiration Use Efficiency, TissueNa and TissueK for sodium and potassium ion accumulation in the shoot tissue respectively, AREA for projected shoot area, MRENDVI for Modification Red Edge Normalized Difference Vegetation Index, PRI for Photochemical Reflectance Index, Temp. for canopy temperature, SFM for shoot fresh mass, MFY for mature fruit yield (g), MFN for mature fruit number, MFM for average mature fruit mass (g), IFY for immature fruit yield (g), IFN for immature fruit number, IFM for average immature fruit mass (g), TYM for total yield mass (g) and TYN for total fruit number. Traits recorded under greenhouse conditions carry prefix TPA (for The Plant Accelerator), whereas traits recorded under field conditions carry prefix KAU (for King Abdulaziz University farm). Traits recorded under control and salt stress conditions are indicated with C or S, respectively, at the end of each trait.

**Supplemental Table S15. Significantly associated SNPs with S.pimpinellifolium traits recorded under greenhouse conditions.** The individual associations are listed within each trait. The traits are abbreviated as: The individual traits are abbreviated with AGR for absolute growth rate, RGR for relative growth rate, TR for transpiration rate, TUE for transpiration use efficiency, TissueK and TissueN for potassium and sodium accumulation, and SFM for shoot fresh mass. Traits recorded under C and S stress conditions are indicated with C or S, respectively, at the end of each trait.

**Supplemental Table S16. The loci of interest identified with the traits recorded in the greenhouse experiment.** Six loci were identified within the GWAS associations identified for S. pimpinellifolium grown under control and salt stress conditions. The traits associated within each of the loci are abbreviated as AGR for absolute growth rate, RGR for relative growth rate, TissueN for sodium accumulation. Traits recorded under C and S stress conditions are indicated with C or S, respectively, at the end of each trait. The genes within the linkage disequilibrium (LD) region of each locus are listed, with their predicted function based on the homology with related species. The position of SNPs in individual genes is reported under "Notes" section.

