## Supplemental Figures for "Deciphering Salt Stress Responses in *Solanum pimpinellifolium* through High-Throughput Phenotyping"

### Supplementary material

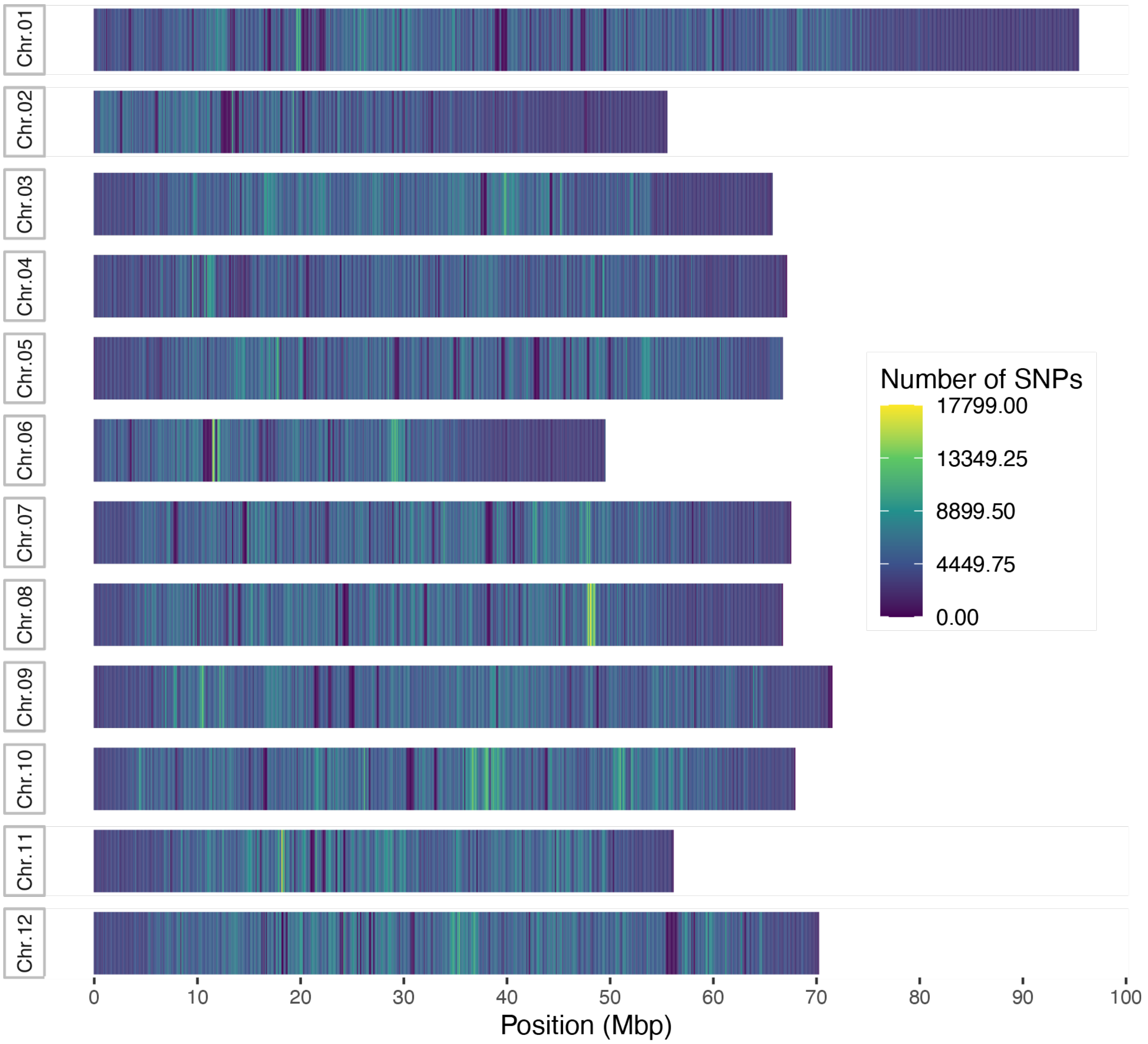


**Figure S1. The density of SNP identified in S. pimpinellifolium diversity panel distributed over 12 chromosomes.** All the mapped SNPs (including heterozygous SNPs) are plotted over their respective position within the *S. pimpinellifolium* calculated in a bin size of 100 kb. The yellow and blue shades represent high and low number of identified SNPs respectively.

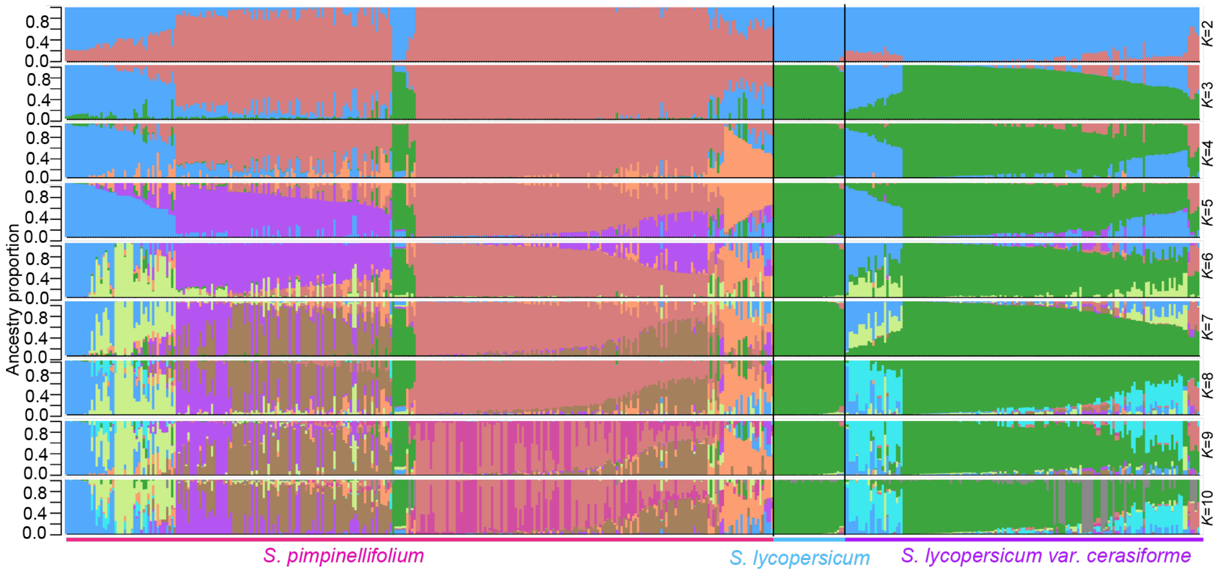


**Figure S2. Population structure of wild and cultivated tomatoes.** Population structure (from *K* = 2 to *K* = 10) of 482 *S. pimpinellifolium*, *S. lycopersicum* and *S. lycopersicum var. cerasiforme* samples estimated with sNMF. Each bar represents a sample and the bars are filled by colors representing the likelihood of membership to each ancestry.

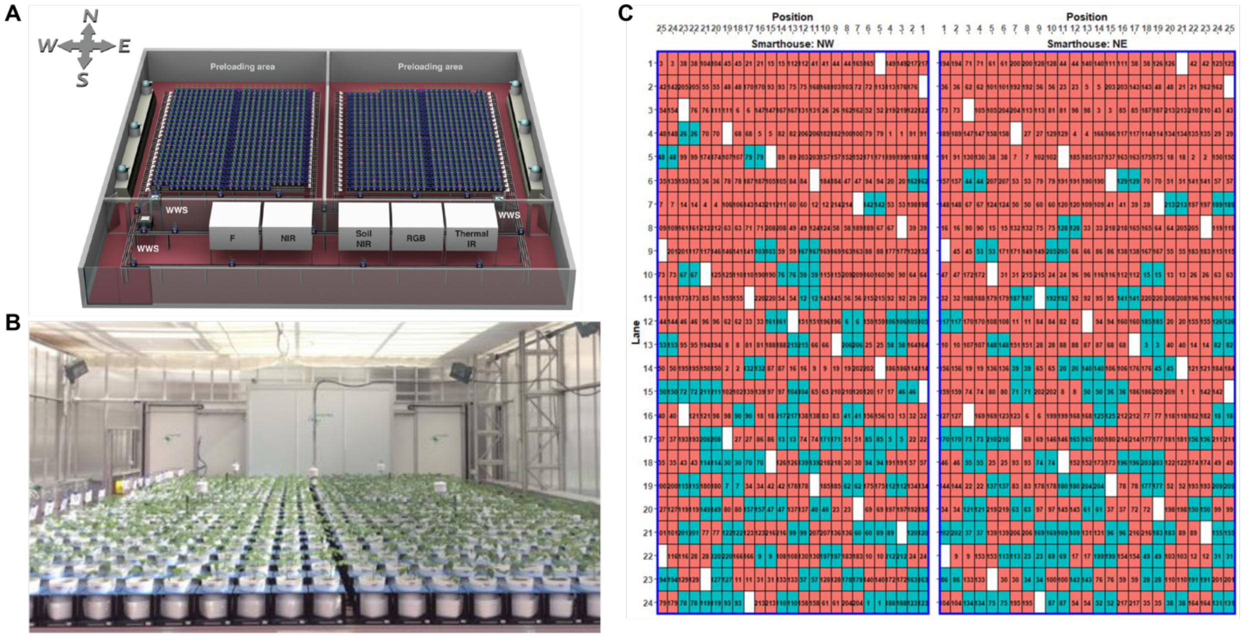


**Figure S3. Design of the greenhouse in The Plant Accelerator, for controlled environment phenotyping. A)** Model of the two TPA® smarthouses and phenotyping stations used in this experiment. WWS: weighing and watering station, Thermal IR: thermal infrared imaging station*, RGB: Red-Blue-Green imaging station, NIR: near infrared imaging*; F: fluorescence imaging*. (*) Imaging stations were not functional for this experiment. Image credit: Heno Hwang. **B)** *S. pimpinellifolium* plants growing in the NW smarthouse. **C)** Plot of the design showing the allocation of accessions to main plots, each main plot being comprised of two consecutive carts. The two salt treatment were randomized to the two consecutive carts within a main plot. The accessions are numbered from 1 to 220. The cells coloured orange are a complete replicate the 220 accessions within a greenhouse, and the cells coloured blue are the second replicates of the 68 accessions that had two replicates within a greenhouse. The white cells are the evaporation pots.

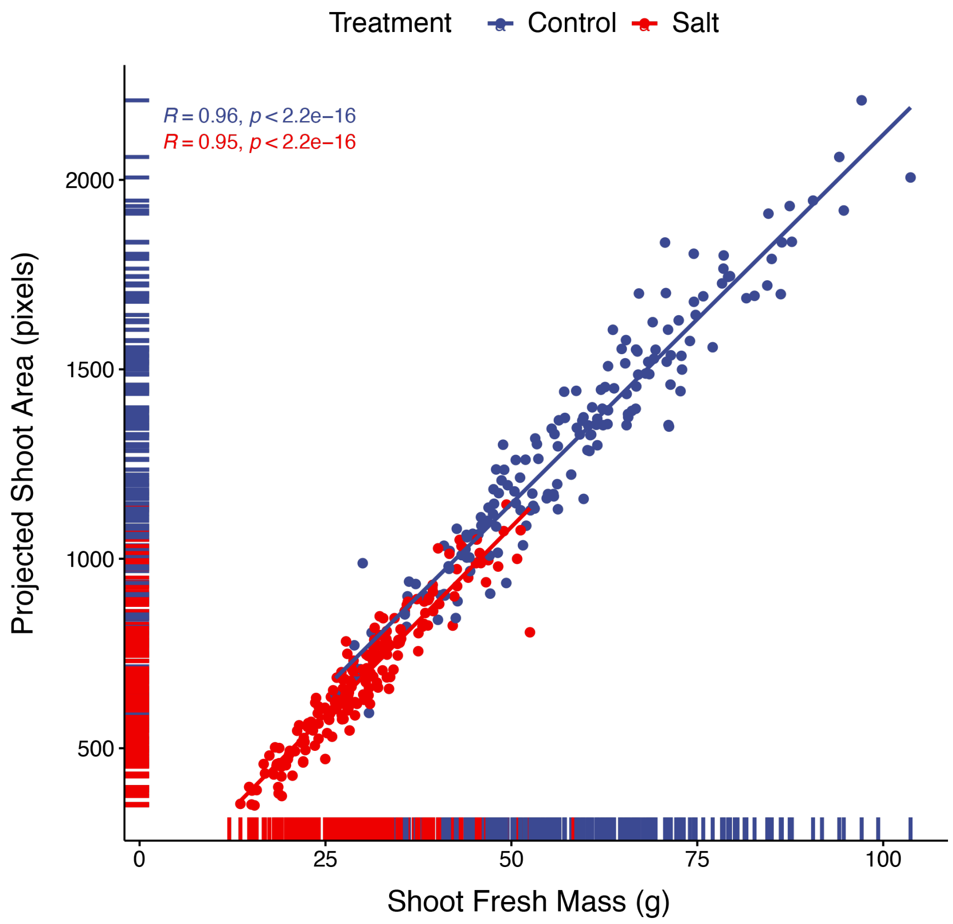


**Figure S4. The relationship between shoot fresh mass and projected shoot area in *S. pimpinellifolium* plants grown under environmentally controlled conditions.** The 220 *S. pimpinellifolium* accessions were grown under greenhouse conditions with and without exposure to salt stress for 2 weeks. The shoot fresh weight, recorded at the last day of the experiment, was compared to projected shoot area, recorded at the same day. The Spearman correlation was calculated for each treatment, together with the correlation p-value. The individual points represent genotype mean at individual condition. The lines represent the linear regression between the two traits.

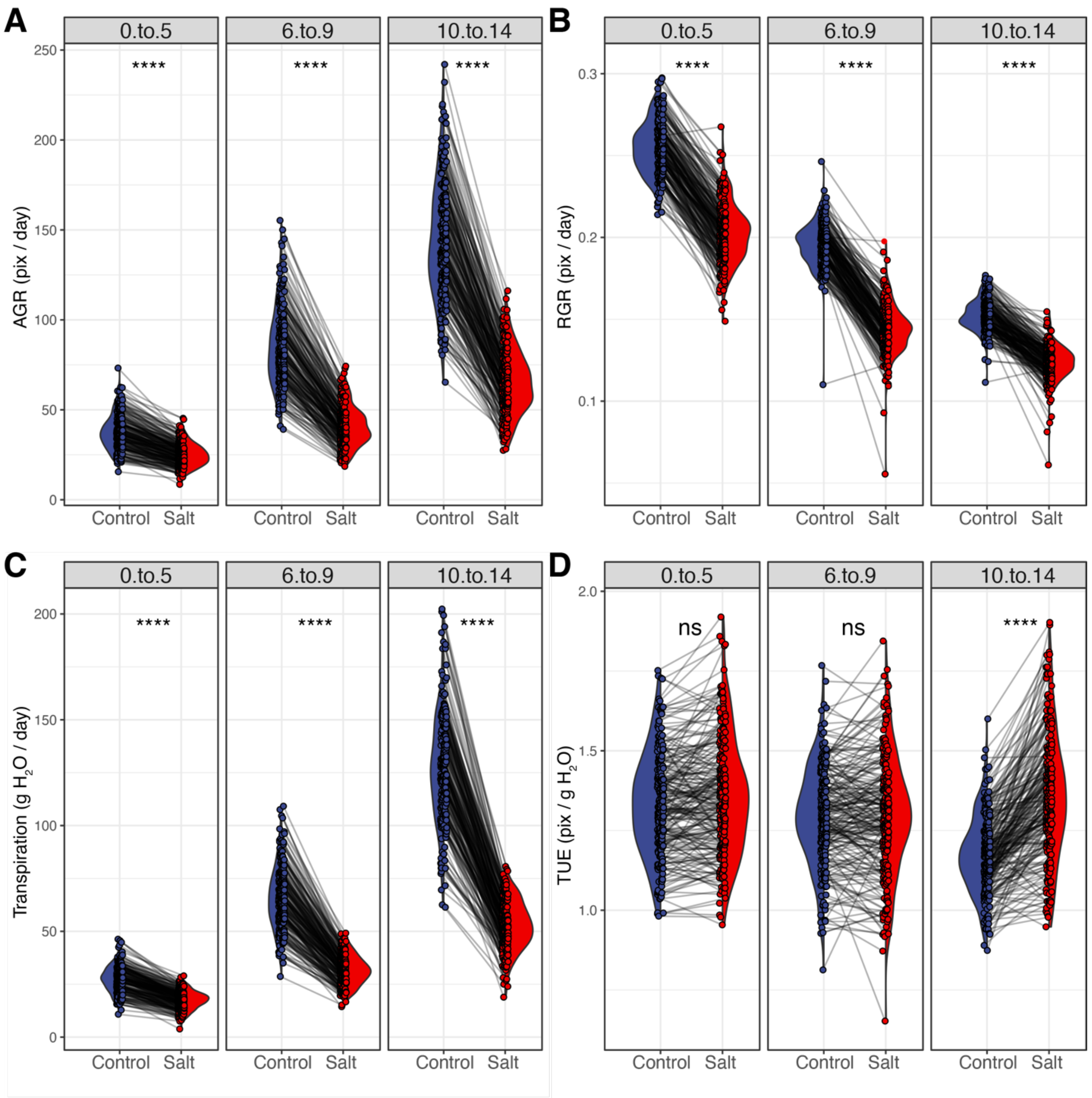


**Figure S5. Data complexity was reduced by summarizing plant growth and transpiration rates into three intervals.** The change in the trait value observed over the course of two weeks of Control or Salt stress treatment (Figure 2), was summarized by calculating a genotypic mean values for three intervals: 0 to 5 days, 6 to 9 days, and 10 to 14 days. We examined stress-induced changes in **A)** Absolute Growth Rate (AGR), **B)** Relative Growth Rate (RGR), **C)** Transpiration Rate and **D)** Transpiration Use Efficiency (TUE). The individual lines in describe the change within a genotype observed between the treatments. The differences between treatments were tested using one-way ANOVA, and *, **, *** and **** indicate p-values below 0.05, 0.01, 0.001 and 0.0001 respectively.

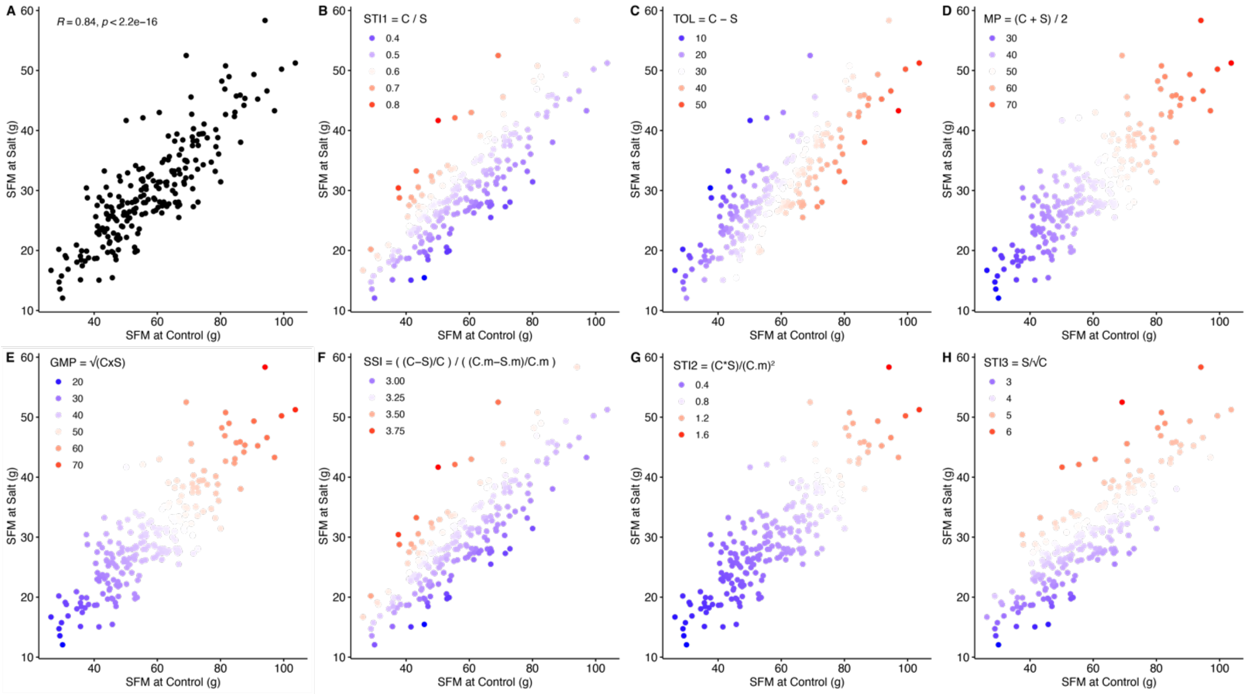


**Figure S6. Evaluation of the *S.pimpinellifolium* accessions and identification of the most tolerant genotypes based on shoot fresh weight of plants grown under controlled environmental conditions.** The performance of *S. pimpinellifolium* accessions was evaluated based on the Shoot Fresh Mass (SFM) accumulated at the end of experiment (2 weeks after initial Control / Salt stress treatment). **A)** The relation between SFM of each genotype under Control and Salt stress conditions was examined using scatter plot, and the Spearman correlation coefficient (R) was calculated, along with the correlation p-value. Subsequently, we highlighted the accessions that showed highest and lowest performance based on the 7 stress tolerance indices: **B)** Salt tolerance index (STI = Salt / Control), **C)** Tolerance index (TOL = Control - Salt), **D)** Mean productivity index (MP = (Control + Salt) / 2), **E)** Geometric Mean Productivity (GMP = square root of (Control x Salt), **F)** Stress Susceptibility Index (SSI = ((Control - Salt) / Control)/ ((population mean Control - population mean Satl) / population mean Control), **G)** Stress Tolerance Index (STI2 = (Control x Salt) / population mean Control$^2$), **H)** Stress Weighted Performance Index (SWP = Salt / square root of Control). The color scale for each index is contained within each graph, with red and blue points representing the best and poorest performing genotypes, respectively.

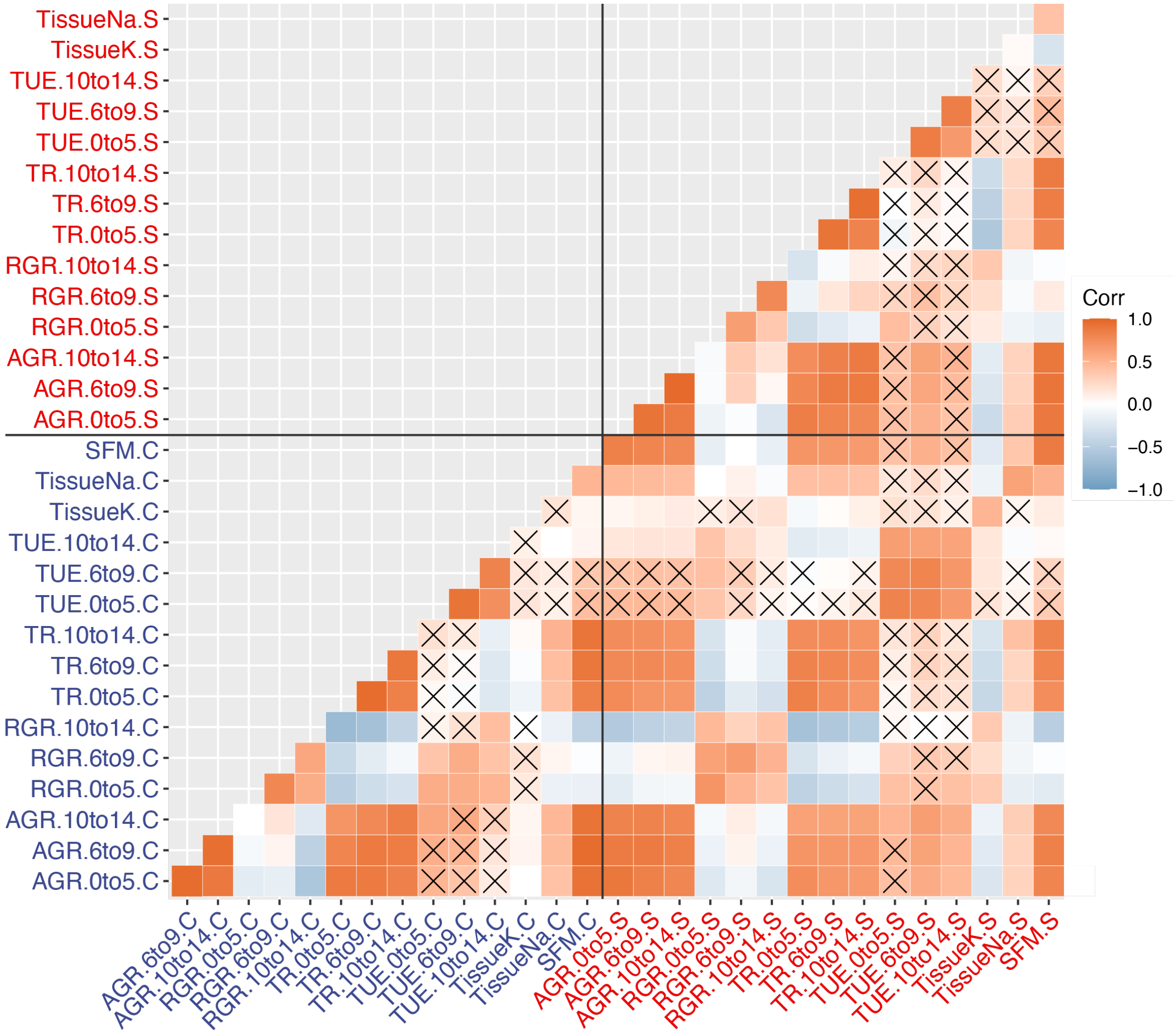


**Figure S7. Correlations between the traits recorded in the greenhouse experiment.** The pearson's correlation coefficients were calculated between each combination of traits. Positive and negative correlations are depicted using orange and blue hues, respectively. Non-significant correlations (p-value > 0.05) are indicated with an X. The individual traits are abbreviated with AGR for absolute growth rate, RGR for relative growth rate, TR for transpiration rate, TUE for transpiration use efficiency, TissueK and TissueN for potassium and sodium accumulation, and SFM for shoot fresh mass. Traits recorded under control and salt stress conditions are indicated with C or S, respectively, at the end of each trait.

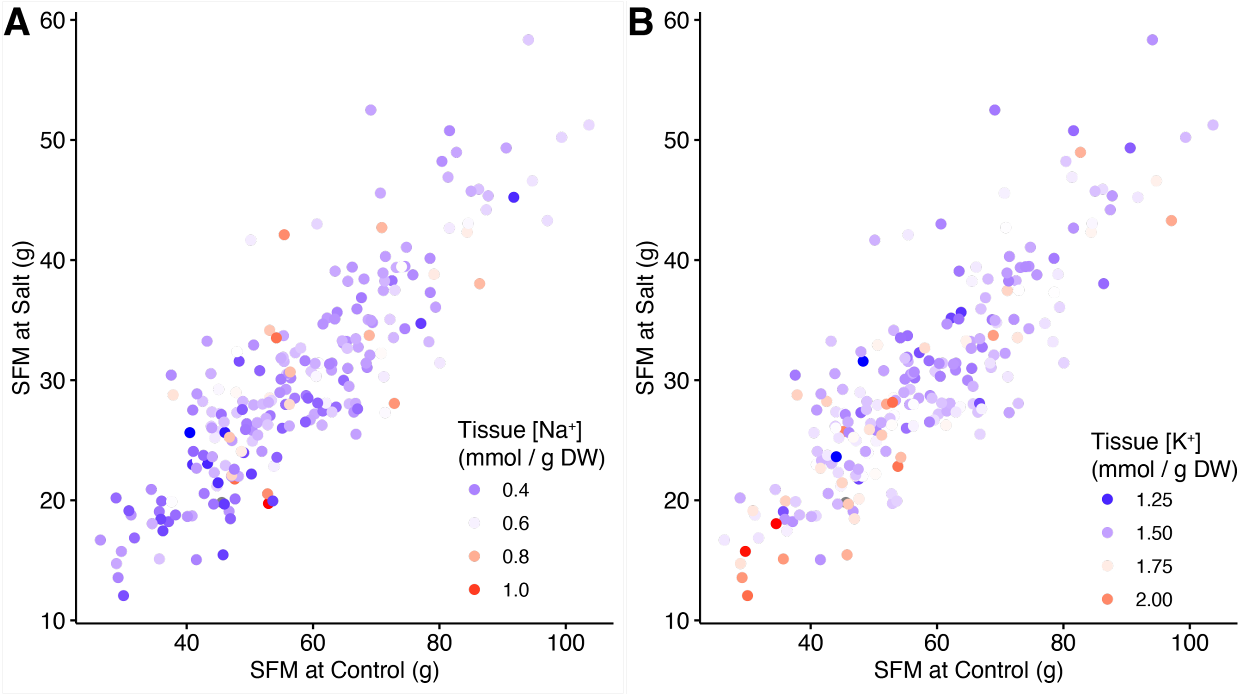


**Figure S8. Evaluation of the *S.pimpinellifolium* accessions for sodium and potassium accumulation.** The performance of 220 *S. pimpinellifolium* accessions was evaluated based on the shoot fresh mass (SFM) accumulated at the end of experiment (2 weeks after initial Control / Salt stress treatment). **A)** The relation between SFM of each genotype under Control and Salt stress conditions was examined using scatter plot, and the A) leaf sodium ($Na^+$) and B) potassium ($K^+$) accumulation is superimposed on the plant performance using the color scale. Color scale for each element is included within each plot, with red and blue points representing high and low ion accumulation respectively.

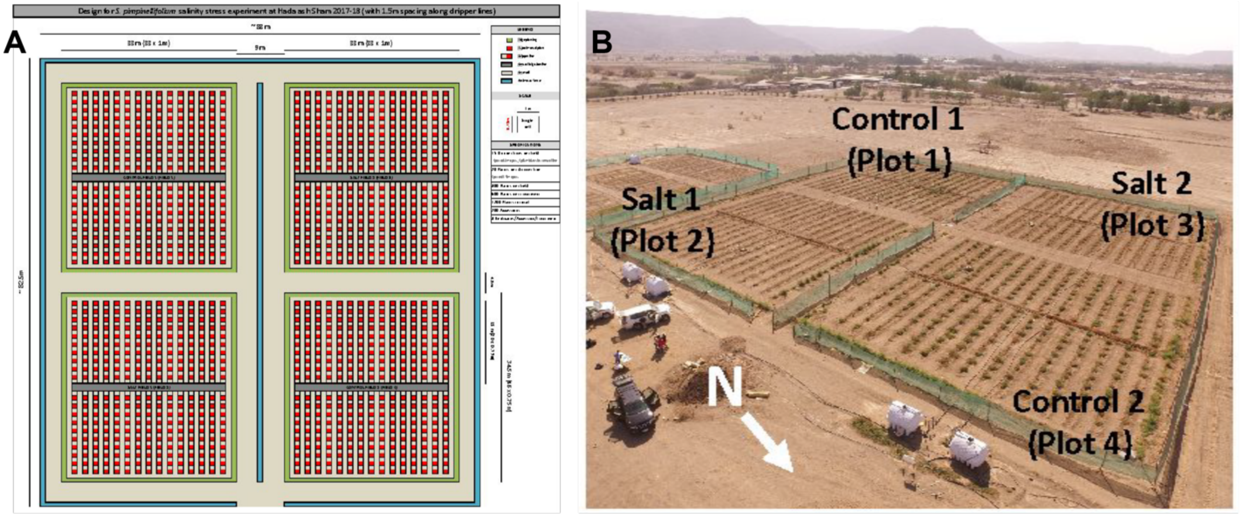


**Figure S9. Design of the field experiment. A)** Layout of field site **B)** Aerial view of the field site, with different plots labelled and due north indicated.

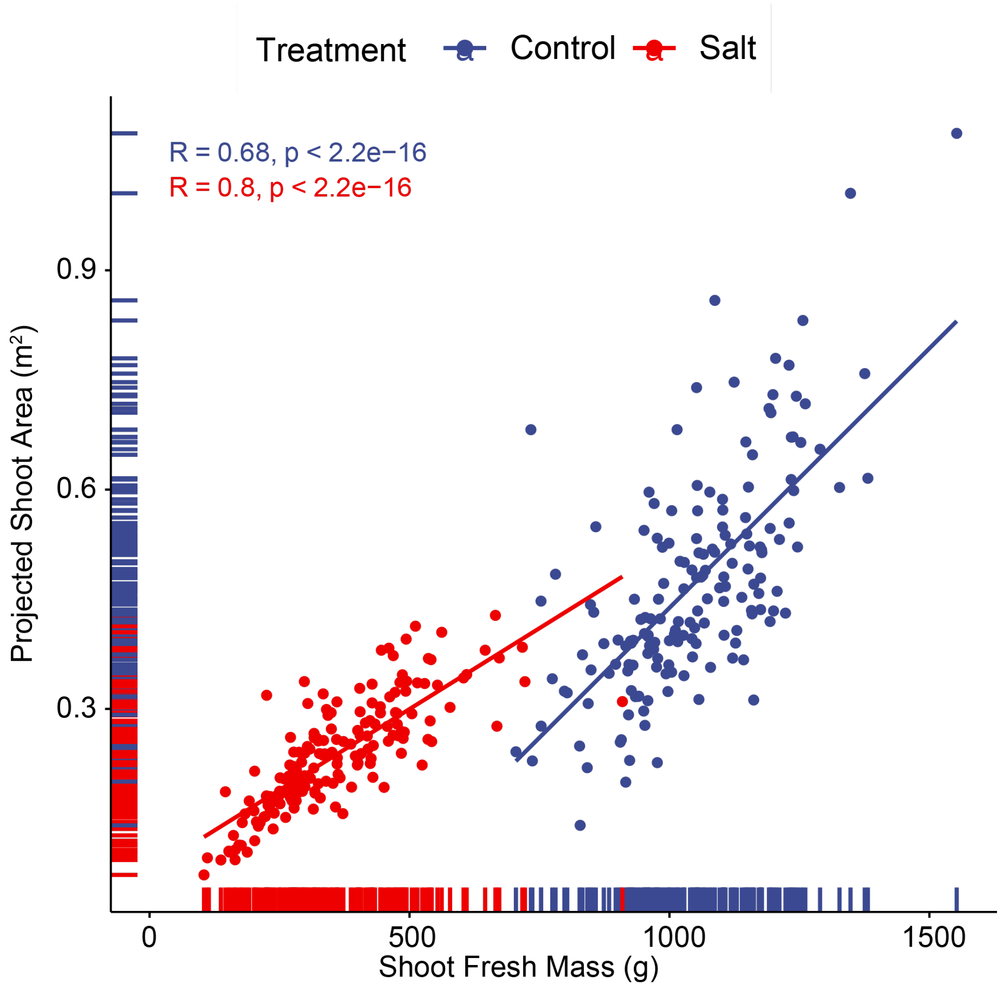


**Figure S10. The relationship between shoot fresh mass and projected shoot area in *S. pimpinellifolium* plants grown under field conditions.** The 119 *S. pimpinellifolium* accessions were grown under field conditions with and without exposure to salt stress for 11 weeks. The shoot fresh weight, recorded at the last day of the experiment, was compared to projected shoot area, recorded at the same day. The Spearman correlation was calculated for each treatment, together with the correlation p-value. The individual points represent genotype mean at individual condition. The lines represent the linear regression between the two traits.

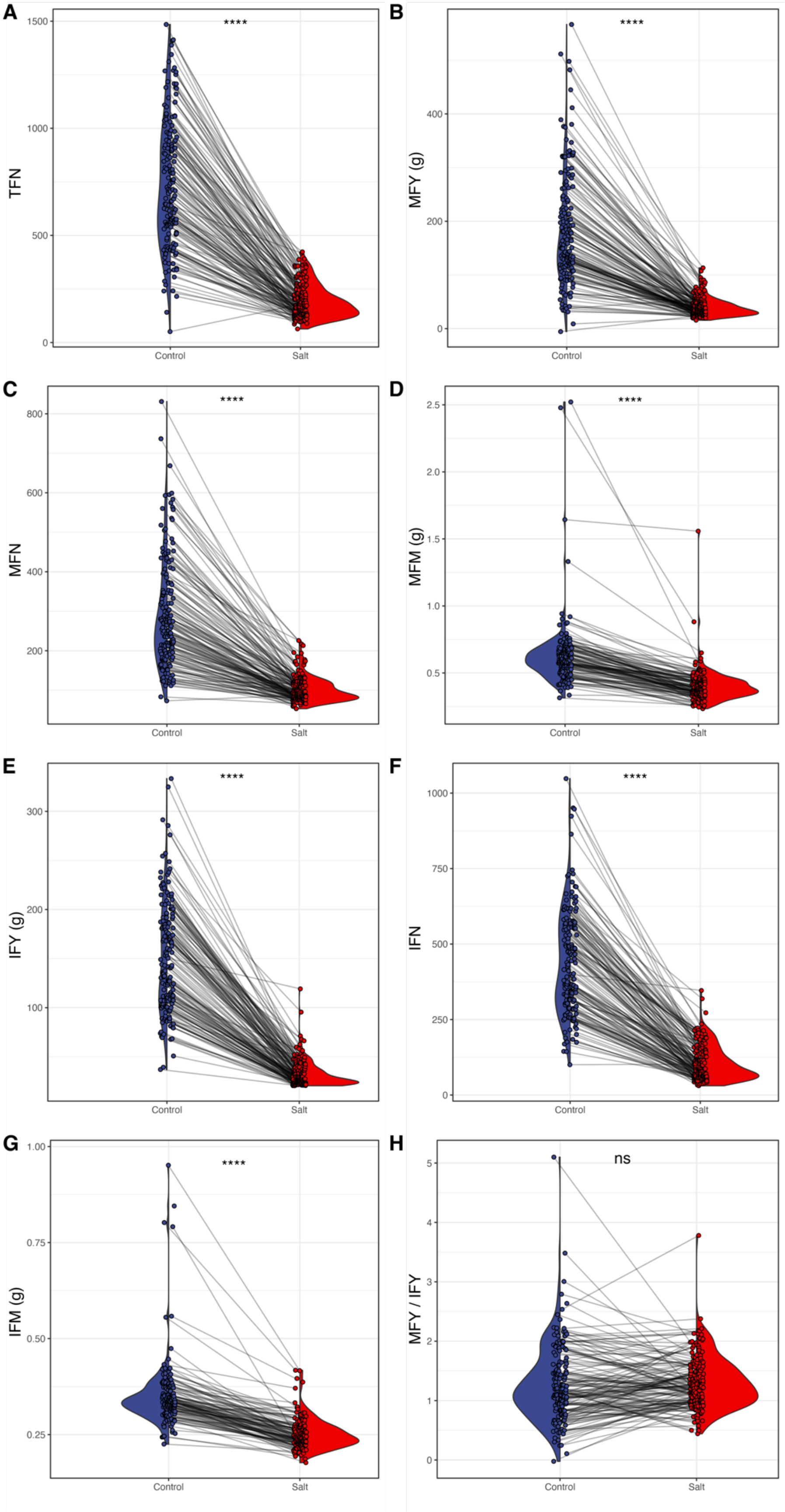


**Figure S11. Salt treatment reduced plant productivity, but not fruit maturation.** The *S. pimpinellifolium* plants exposed to control or salt stress treatment for over 60 days were manually harvested, and evaluated for **A)** total fruit number (TFN) **B)** mature fruit yield (g), **C)** Mature fruit number (MFN), **D)** Mature fruit mass (g), **E)** Immature Fruit Yield (g), **F)** Immature fruit number (IFN), **G)** Immature fruit mass (g) and **H)** ratio of mature to immature fruit yield (MFY / IFY). The individual lines in describe the change within a genotype observed between the treatments. The differences between Control and Salt stress treatments were tested using ANOVA, and *, **, *** and **** indicate p-values below 0.05, 0.01, 0.001 and 0.0001 respectively.

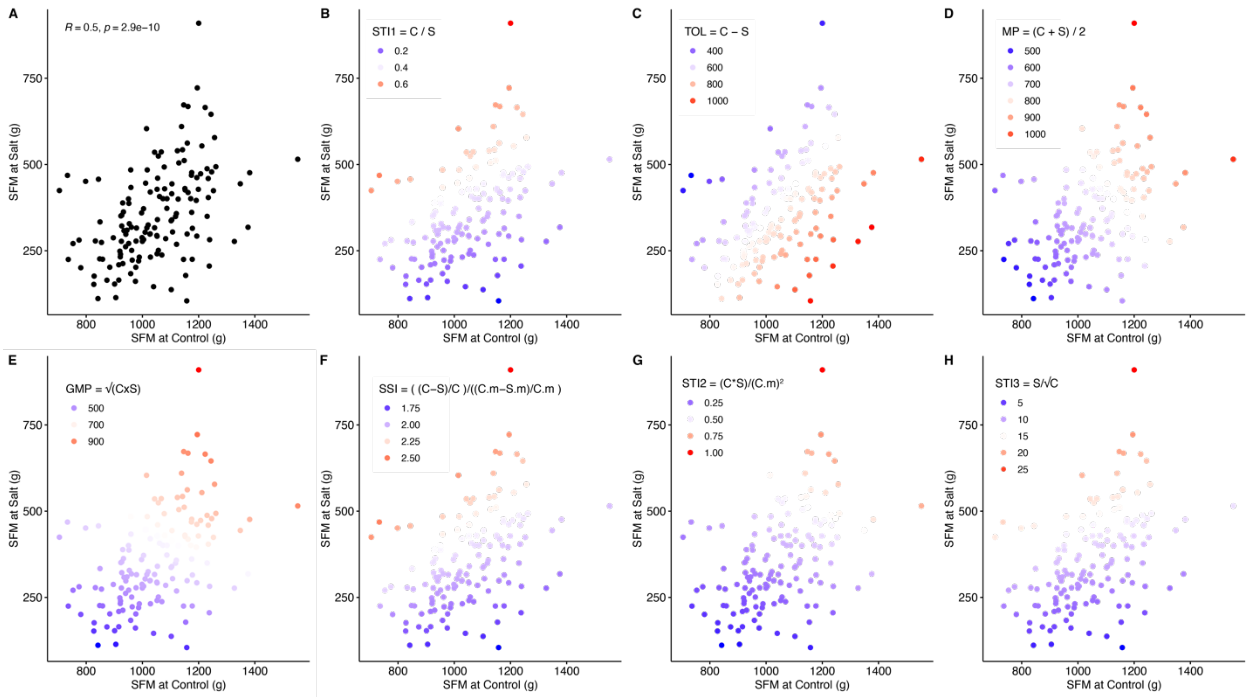


**Figure S12. Evaluation of the *S.pimpinellifolium* accessions and identification of the most tolerant genotypes based on shoot fresh weight of plants grown under field conditions.** The performance of 199 *S. pimpinellifolium* accessions was evaluated based on the shoot fresh mass (SFM) accumulated at the end of experiment (11 weeks after initial Control / Salt stress treatment). **A)** The relation between SFM of each genotype under Control and Salt stress conditions was examined using scatter plot, and the Spearman correlation coefficient (R) was calculated, along with the correlation p-value. Subsequently, we highlighted the accessions that showed highest and lowest performance based on the 7 stress tolerance indices: **B)** Salt tolerance index (STI = Salt / Control), **C)** Tolerance index (TOL = Control - Salt), D) Mean productivity index (MP = (Control + Salt) / 2), **E)** Geometric mean productivity (GMP = square root of (Control x Salt), **F)** Stress Susceptibility Index (SSI = ((Control - Salt) / Control)/ ((population mean Control - population mean Satl) / population mean Control), **G)** Stress Tolerance Index (STI2 = (Control x Salt) / population mean Control$^2$), **H)** Stress Weighted Performance Index (SWP = Salt / square root of Control). The color scale for each index is contained within each graph, with red and blue points representing the best and poorest performing genotypes.

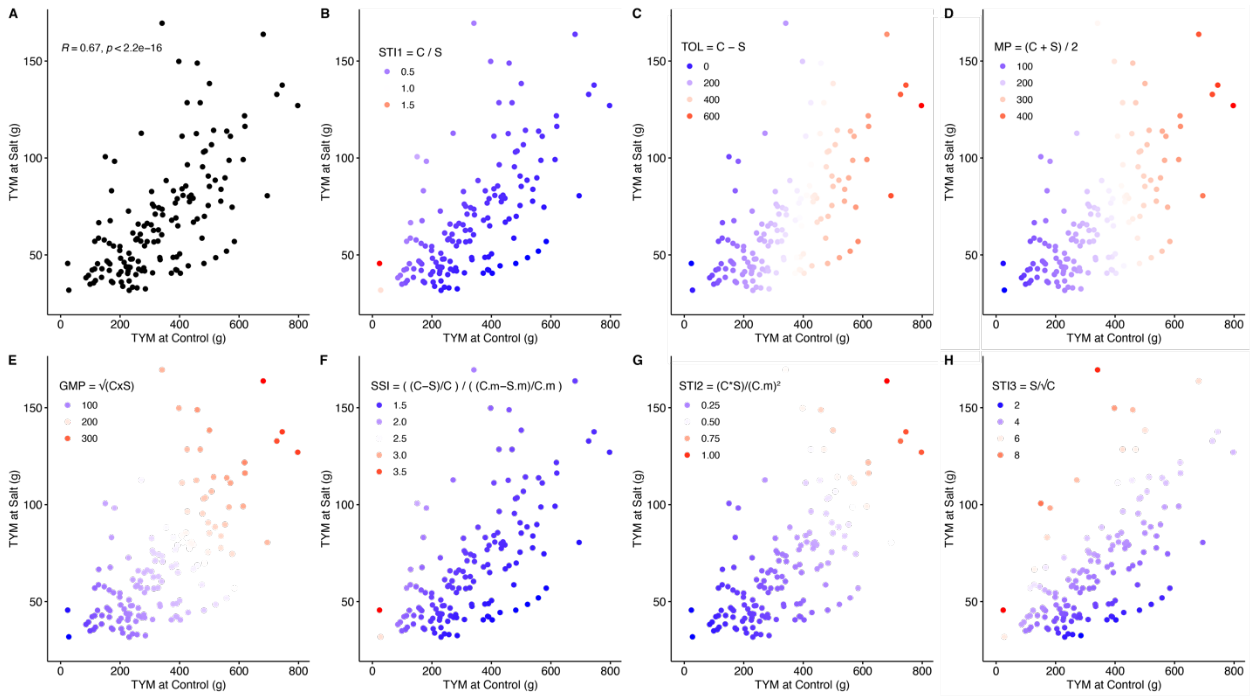


**Figure S13. Evaluation of the *S.pimpinellifolium* accessions and identification of the most tolerant genotypes based on total yield mass of plants grown under field conditions.** The performance of 199 *S. pimpinellifolium* accessions was evaluated based on the total yield mass (TYM) accumulated at the end of experiment (11 weeks after initial Control / Salt stress treatment). **A)** The relation between TYM of each genotype under Control and Salt stress conditions was examined using scatter plot, and the Spearman correlation coefficient (R) was calculated, along with the correlation p-value. Subsequently, we highlighted the accessions that showed highest and lowest performance based on the 7 stress tolerance indices: **B)** Salt tolerance index (STI = Salt / Control), **C)** Tolerance index (TOL = Control - Salt), D) Mean productivity index (MP = (Control + Salt) / 2), **E)** Geometric mean productivity (GMP = square root of (Control x Salt), **F)** Stress Susceptibility Index (SSI = ((Control - Salt) / Control)/ ((population mean Control - population mean Satl) / population mean Control), **G)** Stress Tolerance Index (STI2 = (Control x Salt) / population mean Control$^{2}$), **H)** Stress Weighted Performance Index (SWP = Salt / square root of Control). The color scale for each index is contained within each graph, with red and blue points representing the best and poorest performing genotypes.

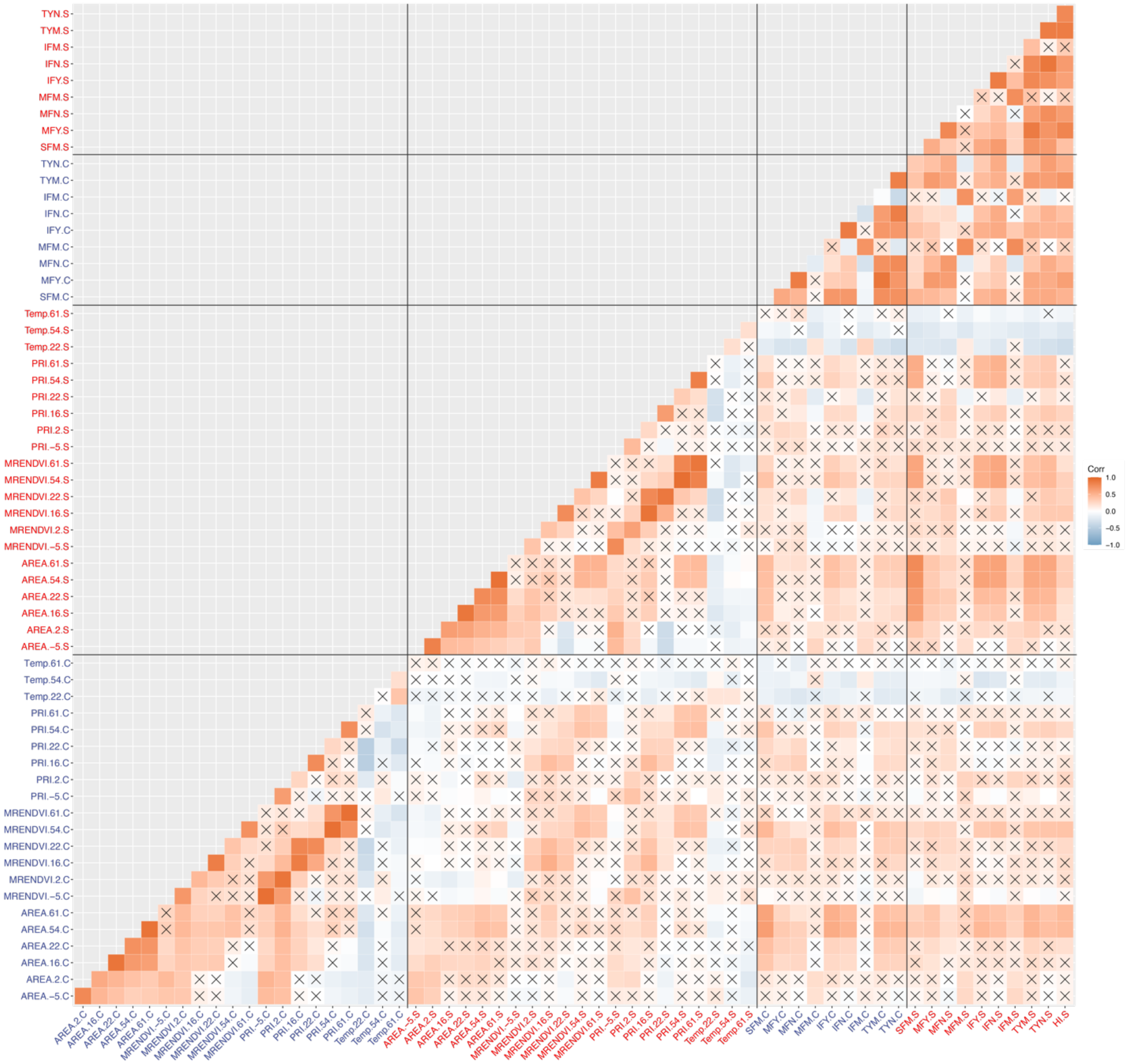


**Figure S14. Correlations between the traits recorded in the field experiment.** The pearson's correlation coefficients were calculated between each combination of traits. Positive and negative correlations are depicted using orange and blue hues, respectively. Non-significant correlations (p-value > 0.05) are indicated with an X. The individual traits are abbreviated with AREA for projected shoot area, MRENDVI for Modification Red Edge Normalised Difference Vegetation Index, PRI for Photochemical Reflectance Index, Temp. for canopy temperature, SFM for shoot fresh mass, MFY for mature fruit yield (g), MFN for mature fruit number, MFM for average mature fruit mass (g), IFY for immature fruit yield (g), IFN for immature fruit number, IFM for average immature fruit mass (g), TYM for total yield mass (g) and TYN for total fruit number. Traits recorded at various time points are indicated with a number following the trait description, corresponding to the days after salt stress imposition. Traits recorded under control and salt stress conditions are indicated with C or S, respectively, at the end of each trait.

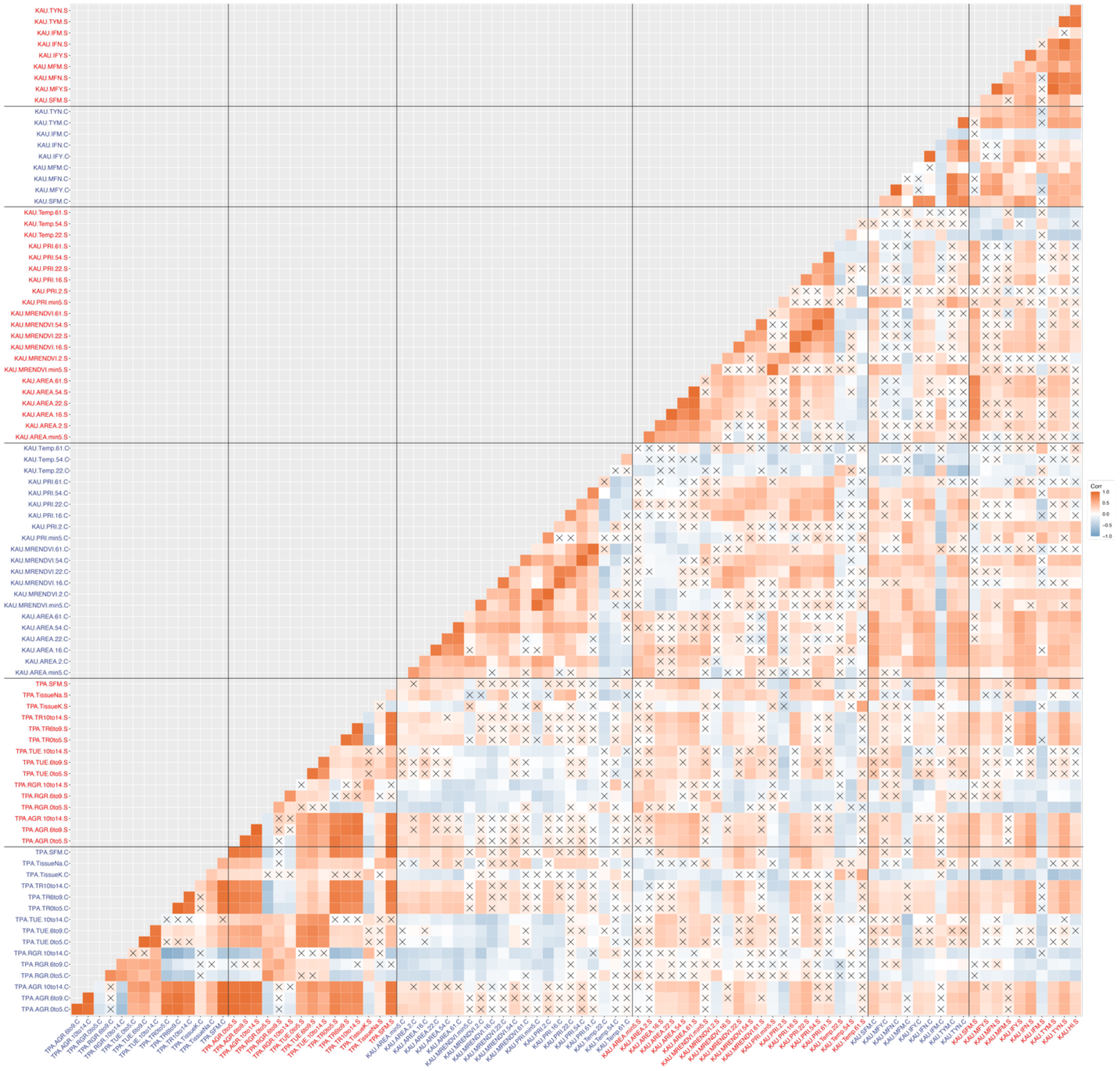

**Figure S15. Correlations between the traits recorded in the greenhouse and field experiment.** The pearson's correlation coefficients were calculated between each combination of traits. Positive and negative correlations are depicted using orange and blue hues, respectively. Non-significant correlations (p-value > 0.05) are indicated with an X. The individual traits are abbreviated with AGR for absolute growth rate, RGR for relative growth rate, TR for transpiration rate, TUE for transpiration use efficiency, TissueK and TissueN for potassium and sodium accumulation, and SFM for shoot fresh mass, AREA for projected shoot area, MRENDVI for Modification Red Edge Normalised Difference Vegetation Index, PRI for Photochemical Reflectance Index, Temp. for canopy temperature, SFM for shoot fresh mass, MFY for mature fruit yield (g), MFN for mature fruit number, MFM for average mature fruit mass (g), IFY for immature fruit yield (g), IFN for immature fruit number, IFM for average immature fruit mass (g), TYM for total yield mass (g) and TYN for total fruit number. Traits recorded at various time points are indicated with a number following the trait description, corresponding to the days after salt stress imposition. Traits recorded under greenhouse conditions carry prefix TPA (for The Plant Accelerator), whereas traits recorded under field conditions carry prefix KAU (for King Abdulaziz University farm). Traits recorded under control and salt stress conditions are indicated with C or S, respectively, at the end of each trait.

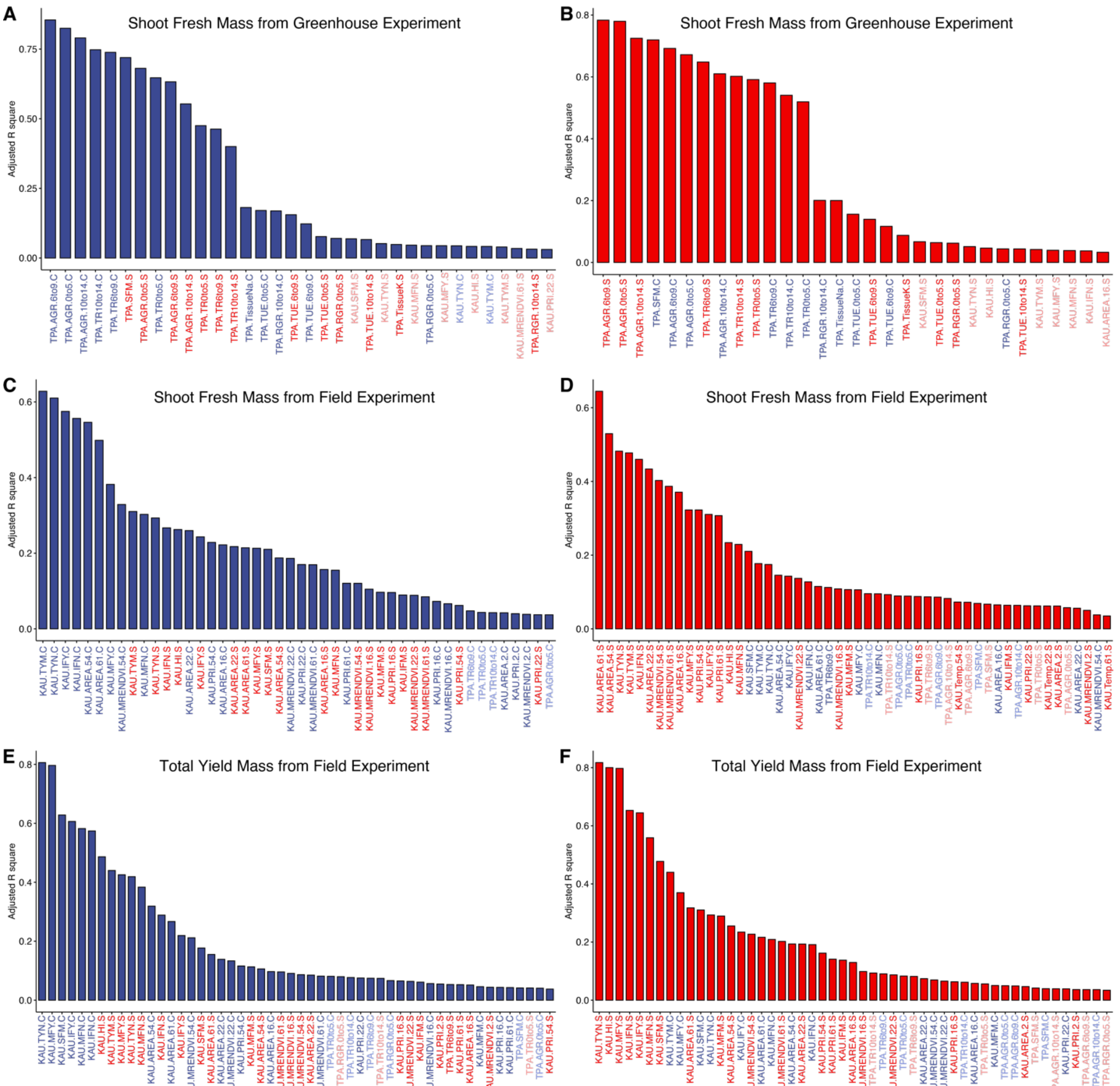


**Figure S16. The regression analysis of plant performance under greenhouse and field conditions.** All of the collected phenotypes over time of the greenhouse (TPA) and field (KAU) experiment were used to perform the linear regression modeling to explain **A)** Shoot fresh mass under control and **B)** salt stress conditions recorded under greenhouse conditions; **C)** shoot fresh mass under control and **D)** salt stress conditions and **E)** total yield mass under control and **F)** salt stress conditions under field conditions. The adjusted regression coefficient (R2) was examined for each measured trait, representing the fraction of explained variation in plant performance. The individual traits are abbreviated with AGR for absolute growth rate, RGR for relative growth rate, TR for transpiration rate, TUE for transpiration use efficiency, TissueK and TissueN for potassium and sodium accumulation, and SFM for shoot fresh mass, AREA for projected shoot area, MRENDVI for Modification Red Edge Normalised Difference Vegetation Index, PRI for Photochemical Reflectance Index, Temp. for canopy temperature, SFM for shoot fresh mass, MFY for mature fruit yield (g), MFN for mature fruit number, MFM for average mature fruit mass (g), IFY for immature fruit yield (g), IFN for immature fruit number, IFM for average immature fruit mass (g), TYM for total yield mass (g) and TYN for total fruit number. Traits recorded under greenhouse conditions carry prefix TPA (for The Plant Accelerator), whereas traits recorded under field conditions carry prefix KAU (for King Abdulaziz University farm). Traits recorded under control and salt stress conditions are indicated with C or S, respectively, at the end of each trait. The traits with not significant (p-value < 0.01) regression coefficients are not displayed.

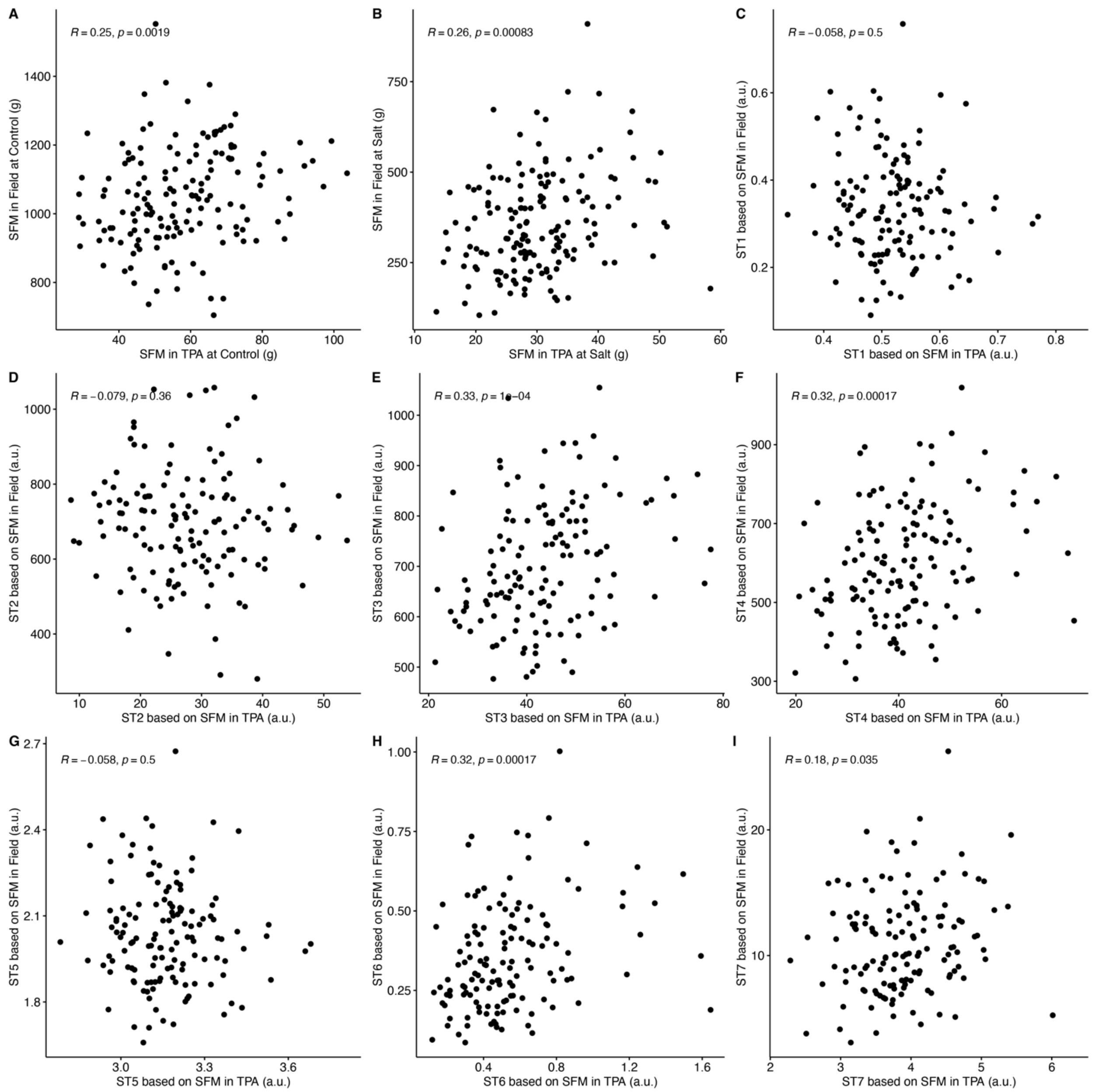


**Figure S17. Evaluation of the *S.pimpinellifolium* accessions and identification of the most tolerant genotypes based on shoot fresh mass of plants grown under controlled environment and field conditions.** The performance of 199 *S. pimpinellifolium* accessions that were included in both controlled environment and field conditions screening was evaluated based on the shoot fresh mass (SFM) accumulated at the end of experiment (2 weeks for controlled conditions, 11 weeks for field experiment). The relation between SFM of each genotype under field and controlled environment experiment at The Plant Accelerator (TPA) was evaluated under **A)** Control and **B)** Salt stress conditions. Additionally, we evaluated the similarities in genotype performance using the various stress indices: **C)** Salt tolerance index (STI = Salt / Control), **D)** Tolerance index (TOL = Control - Salt), **E)** Mean productivity index (MP = (Control + Salt) / 2), **F)** Geometric mean productivity (GMP = square root of (Control x Salt), **G)** Stress Susceptibility Index (SSI = ((Control - Salt) / Control)/ ((population mean Control - population mean Satl) / population mean Control), **H)** Stress Tolerance Index (STI2 = (Control x Salt) / population mean $Control^2$), **I)** Stress Weighted Performance Index (SWP = Salt / square root of Control). Each relationship between control and field conditions was examined using scatter plot, and the Spearman correlation coefficient (R) was calculated, along with the correlation p-value.

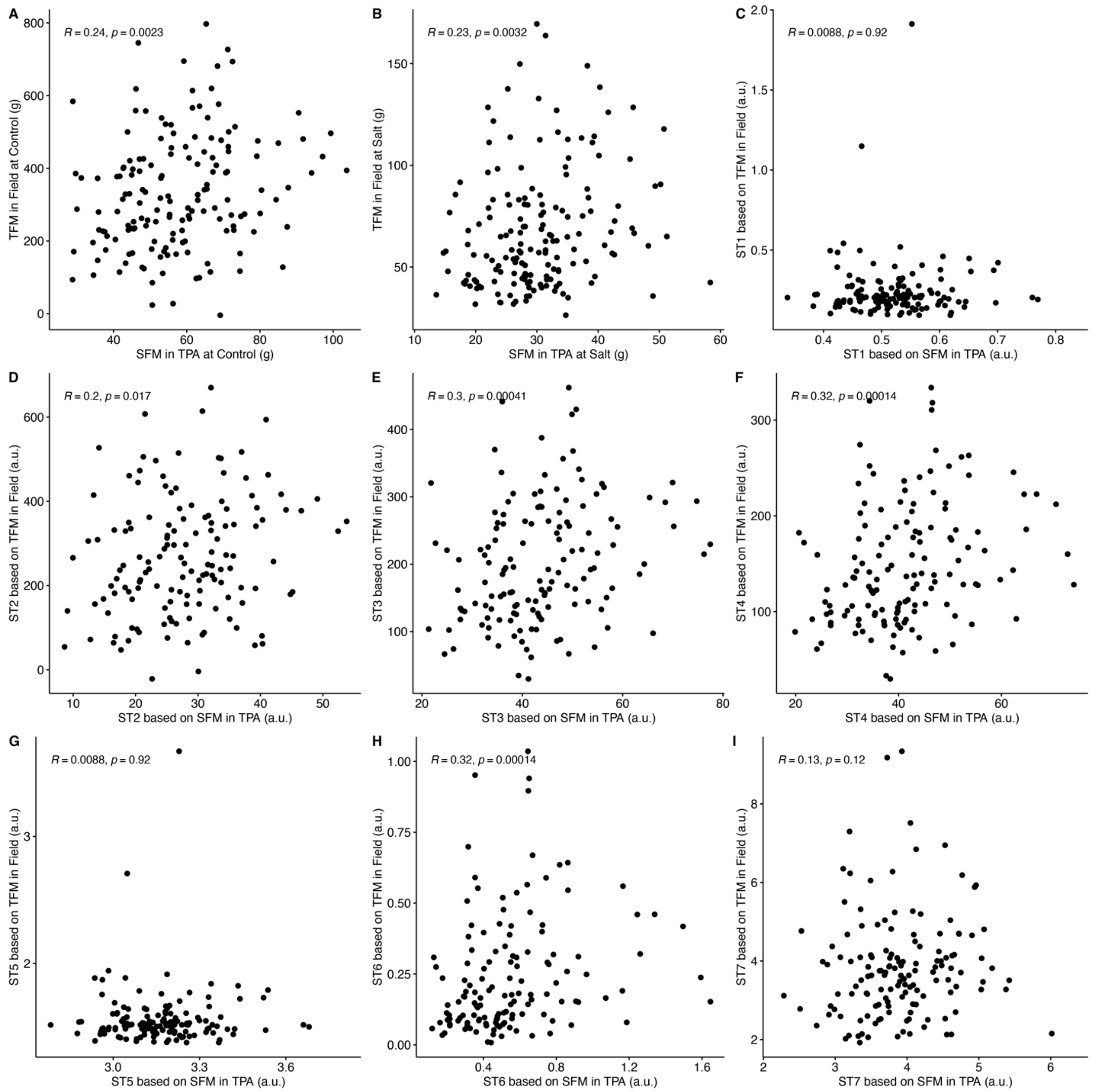


**Figure S18. Evaluation of the *S.pimpinellifolium* accessions and identification of the most tolerant genotypes based on shoot fresh mass of plants grown under controlled environment and total yield mass of plants grown under field conditions.** The performance of 199 *S. pimpinellifolium* accessions that were included in both controlled environment and field conditions screening was evaluated based on the shoot fresh mass (SFM) for controlled environmental conditions and total fruit mass (TFM) for field conditions accumulated at the end of experiment (2 weeks for controlled conditions, 11 weeks for field experiment). The relation between TFM and SFM of each genotype under field and controlled environment experiment at The Plant Accelerator (TPA) was evaluated under **A)** Control and **B)** Salt stress conditions. Additionally, we evaluated the similarities in genotype performance using the various stress indices: **C)** Salt tolerance index (STI = Salt / Control), **D)** Tolerance index (TOL = Control - Salt), **E)** Mean productivity index (MP = (Control + Salt) / 2), **F)** Geometric mean productivity (GMP =

square root of (Control x Salt), **G)** Stress Susceptibility Index (SSI = ((Control - Salt) / Control)/ ((population mean Control - population mean Satl) / population mean Control), **H)** Stress Tolerance Index (STI2 = (Control x Salt) / population mean Control$^2$), **I)** Stress Weighted Performance Index (SWP = Salt / square root of Control). Each relationship between control and field conditions was examined using scatter plot, and the Spearman correlation coefficient (R) was calculated, along with the correlation p-value.

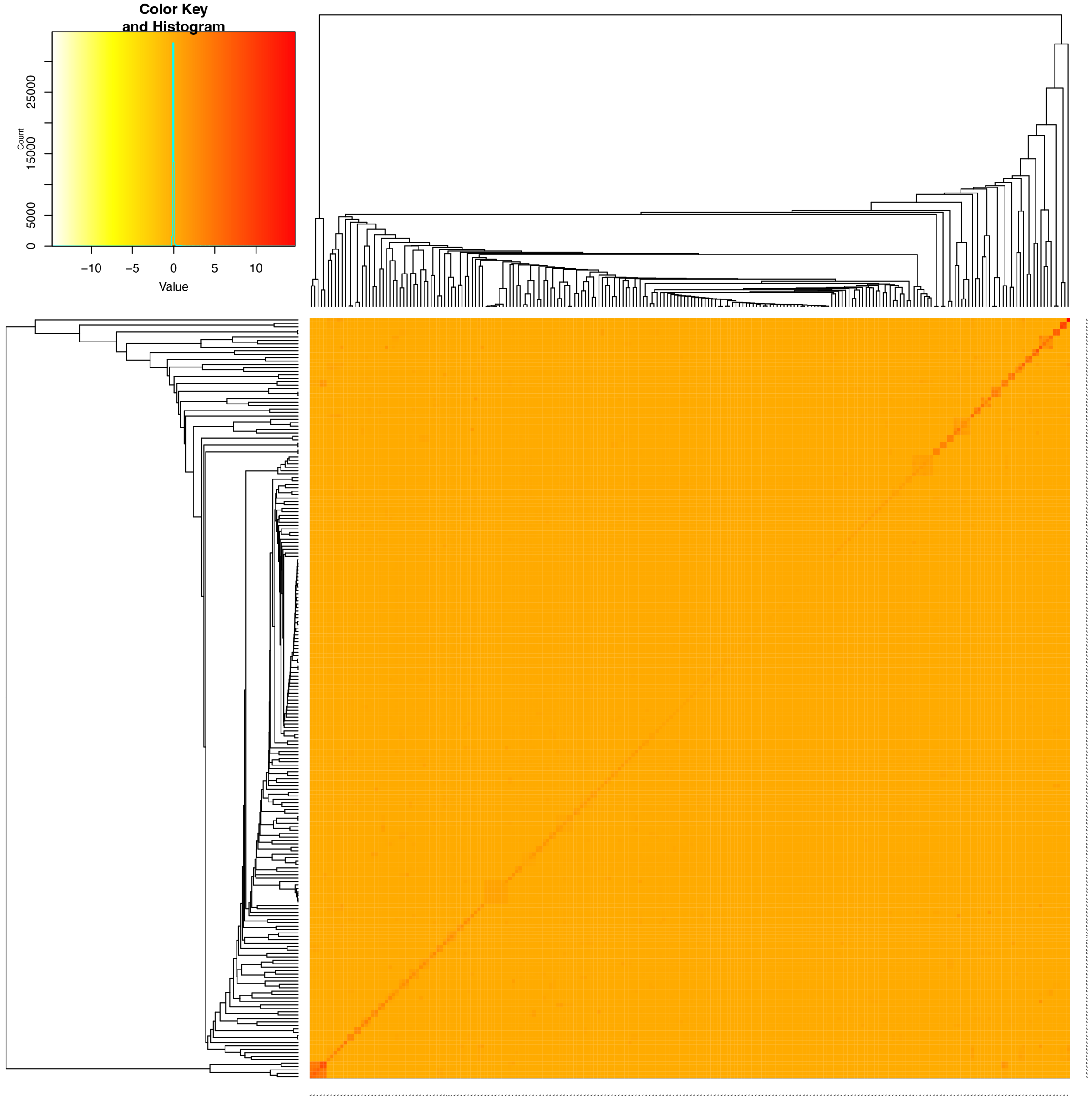


**Figure S19. The kinship matrix for *S. pimpinellifolium* accessions used as a co-factor for Genome Wide Association Study.** The kinship matrix was calculated in GAPIT using 708,545 SNPs that underwent filtering and were subsequently used in GWAS. The kinship between individual accessions is represented as a Z-score value, with red and white hues representing high and low level of kinship. The dendrogram represents hierarchical clustering of 199 genotypes of *S. pimpinellifolium* that were successfully sequenced and used in phenotyping experiments.

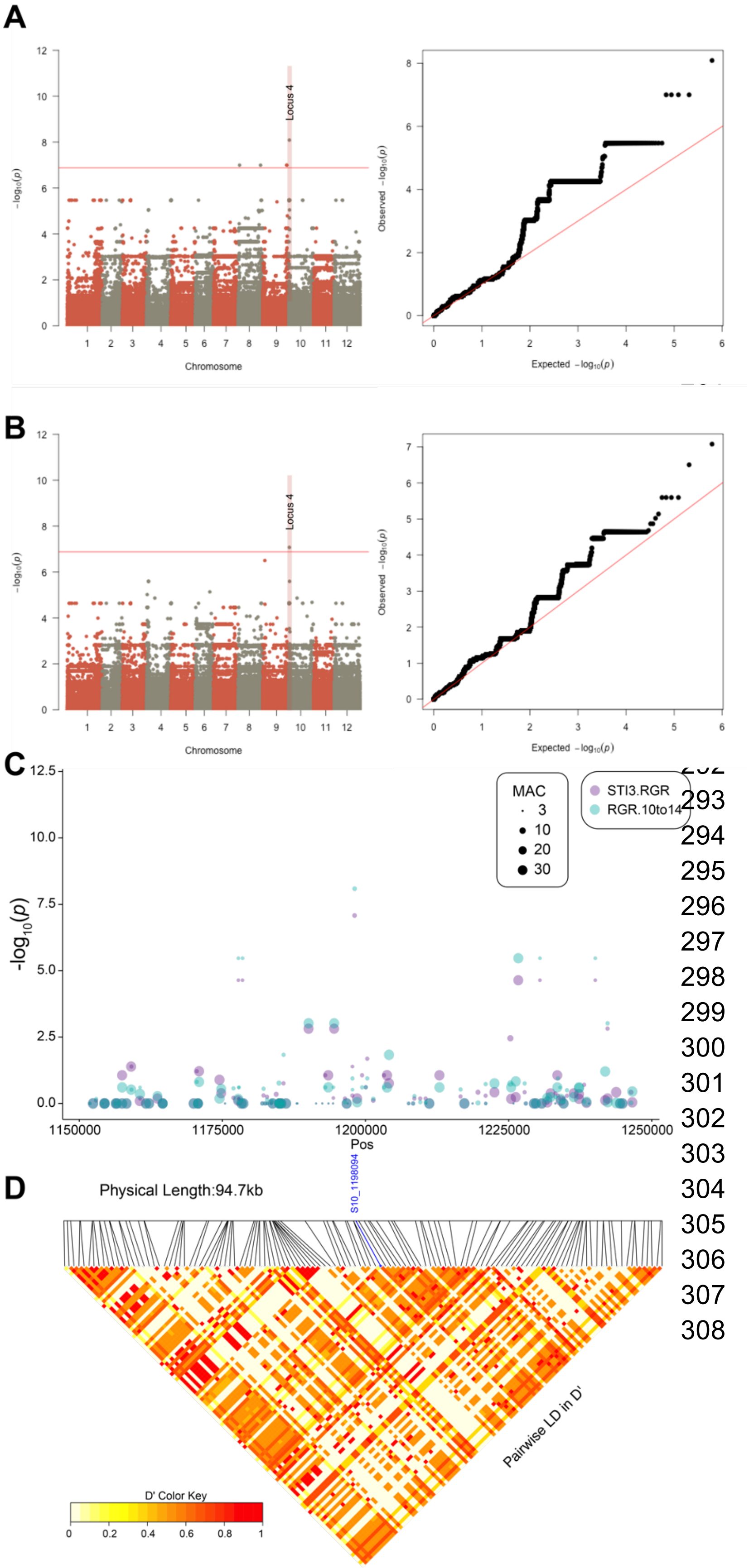


**Figure S20. Natural variation in relative growth rate and salt tolerance index under salt stress in *S. pimpinellifolium* corresponds to one genetic locus.** The Genome Wide Association Study was performed for 220 *S. pimpinellifolim* accessions using 708 545 SNPs. The ASReml GWAS model was applied on **A)** relative growth rate (RGR) in the 3rd interval, and **B)** Salt Tolerance Index (STI = S / C) calculated using RGR for 3rd interval (10 to 14 days after stress application). The identified association consisted of one SNP (position 1198094) on chromosome 9. **C)** The associations in both traits were inspected for other SNPs within the 100 kbp window. **D)** The local linkage disequilibrium (LD) was calculated for all of the SNPs within the 100 kbp window. The significantly associated SNP is highlighted in blue in LD heatmaps, whereas the LD between the individual SNPs is represented as a heatmap, with high intensity of red indicating strong linkage, and yellow hues indicating low linkage.

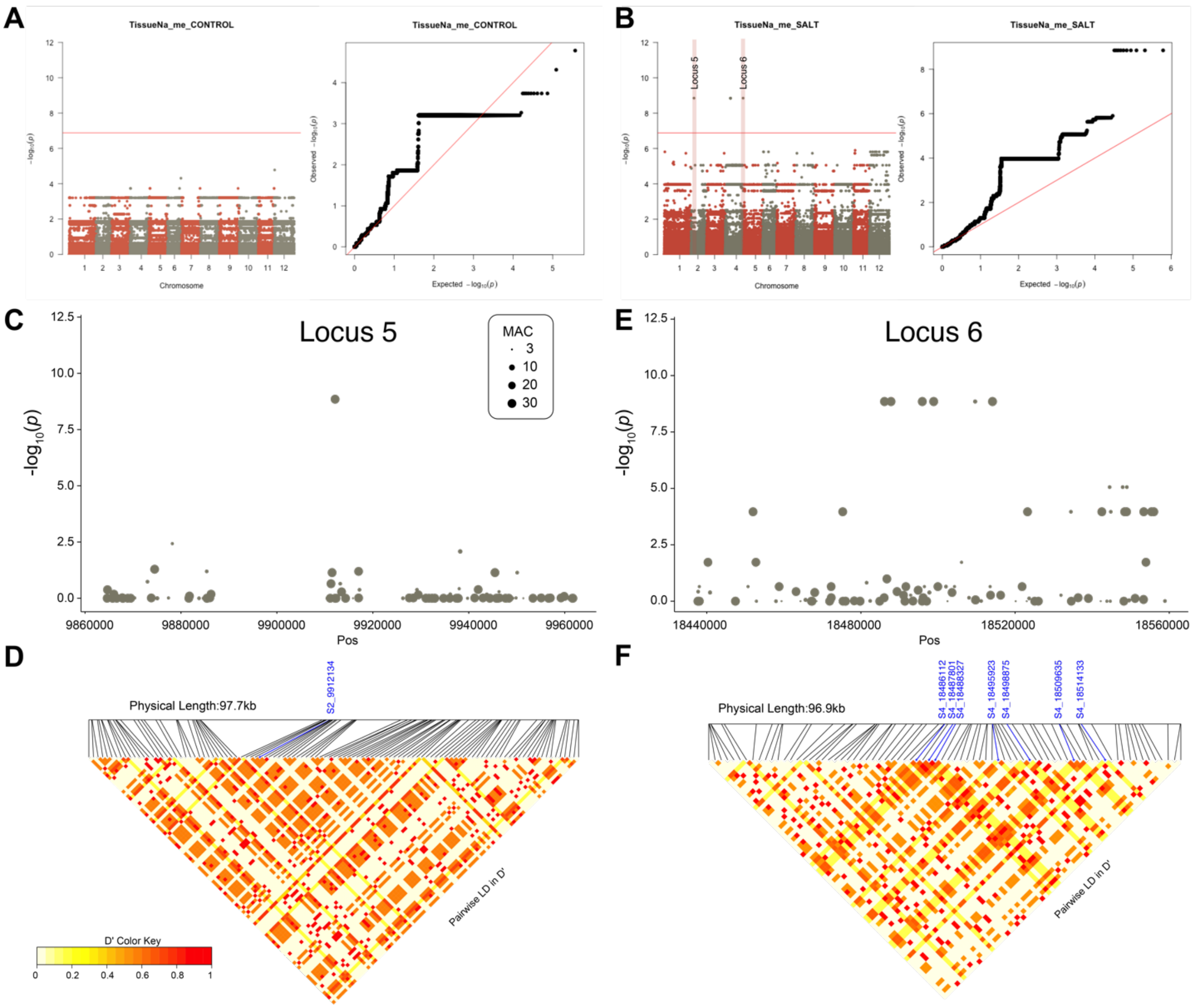


**Figure S21. Natural variation in sodium accumulation in *S. pimpinellifolium* corresponds to genetic variation distributed over two loci.** The Genome Wide Association Study was performed for 220 *S. pimpinellifolim* accessions using 708 545 SNPs. The ASReml GWAS model was applied on sodium ($Na^+$) accumulation in the leaf tissue recorded under **A)** Control and **B)** Salt stress conditions. **C)** The first locus was identified on chromosome 2, position 9912134. The association was inspected for other SNPs within the 100 kbp window. **D)** The local linkage disequilibrium (LD) was calculated for all of the SNPs within the 100 kbp window. **E)** The second locus was identified on chromosome 4, and consisted of seven significantly associated SNPs (positions 18514133, 18509635, 18498875, 18495923, 18488327, 18487801 and 18486112). The region was further inspected for associations within 100 kbp window. F) The local LD was calculated for all of the SNPs within the 100 kbp window. The significantly associated SNPs are highlighted in blue in LD heatmaps, whereas the LD between the individual SNPs is represented as a heatmap, with high intensity of red indicating strong linkage, and yellow hues indicating low linkage.
